# Human CD4^+^CD103^+^ cutaneous resident memory T cells are found in the circulation of healthy subjects

**DOI:** 10.1101/361758

**Authors:** M. M. Klicznik, P. A. Morawski, B. Höllbacher, S. R. Varkhande, S. Motley, L. Kuri-Cervantes, E. Goodwin, M. D. Rosenblum, S. A. Long, G. Brachtl, T. Duhen, M.R. Betts, D. J. Campbell, I. K. Gratz

## Abstract

Tissue-resident memory T cells (T_RM_) persist locally in non-lymphoid tissues where they provide front-line defense against recurring insults. T_RM_ at barrier surfaces express the markers CD103 and/or CD69 which function to retain them in epithelial tissues. In humans, neither the long-term migratory behavior of T_RM_ nor their ability to re-enter the circulation and potentially migrate to distant tissue sites have been investigated. Using tissue explant cultures, we found that CD4^+^CD69^+^CD103^+^ T_RM_ in human skin can downregulate CD69 and exit the tissue.

Additionally, we identified a skin-tropic CD4^+^CD69^−^CD103^+^ population in human lymph and blood that is transcriptionally, functionally and clonally related to the CD4^+^CD69^+^CD103^+^ T_RM_ population in the skin. Using a skin xenograft model, we confirmed that a fraction of the human cutaneous CD4^+^CD103^+^ T_RM_ population can re-enter circulation, and migrate to secondary human skin sites where they re-assume a T_RM_ phenotype. Thus, our data challenge current concepts regarding the strict tissue compartmentalization of CD4^+^ T cell memory in humans.

**One Sentence Summary:** Human CD4^+^CD103^+^ cutaneous resident memory T cells are found in the circulation of healthy subjects, and these cells can seed distant skin sites.

## Introduction

T cell memory is compartmentalized into circulating and tissue-resident cell populations. Whereas circulating memory T cells continually patrol the body via the blood and lymphatics, tissue-resident memory T cell (T_RM_) populations establish residence in non-lymphoid organs, where they can provide potent recall responses *(1)*. T_RM_ populations at barrier surfaces such as the intestines, lungs, and skin are best defined by expression of the markers CD103 and/or CD69, which together function to restrict their recirculation and maintain tissue residence *(2)(3)*. However, despite extensive studies there is no single-cell definition for T_RM_. Instead the term T_RM_ is used to describe a cell population within a tissue that is in substantial disequilibrium with cells in the circulation as measured by depletion, tissue-transplantation, or parabiosis studies *(2)(4)(5)*.

T_RM_ were first identified in the context of CD8^+^ T cell responses to infection *(5)(6).* Although cutaneous CD8^+^ T_RM_ have been well-studied in the mouse, the behavior of CD4^+^ memory T cells in mouse skin has been more controversial, with initial studies demonstrating that CD4^+^ T cells in the skin showed a more dynamic pattern of migration and recirculation than cutaneous CD8^+^ T cells, resulting in their equilibration with the circulating T cell pool *(7)(8).* However, skin inflammation or infection increased recruitment and retention of murine CD4^+^ T cells in the skin *(8)(9),* and in some cases led to the formation of sessile cutaneous CD69^+^CD103^+^ CD4^+^ T cells with superior effector functions *(10)(11)*. Within the skin, T_RM_ are most abundant at the site of initial infection *(11)(12)*. However, long-term maintenance of this biased distribution may pose a disadvantage for a large barrier organ like the skin where pathogen re-encounter at a secondary tissue site is possible.

As in experimental animals, human CD4^+^ T_RM_ are generated in response to cutaneous microbes such as *Candida albicans (11)*, but aberrantly activated or malignant T_RM_ are implicated in skin diseases, including psoriasis and mycosis fungoides *(13)*. However, in studying cutaneous CD4^+^ T_RM_, reliance on animal models can be problematic due to fundamental structural differences in the skin in humans versus mice, and a lack of direct correspondence between cutaneous T cell populations in these species. For instance, whereas nearly all CD4^+^ T cells in murine skin are found in the dermis, the human epidermis is much thicker than in mice, and memory CD4^+^ T cells can be found throughout human skin, in both the dermal and the epidermal compartments *(2).* In human skin, most CD4^+^ T cells express CD69, and a fraction of these are also CD103^+^. Moreover, studies following depletion of circulating T cells with anti-CD52 (alemtuzumab) demonstrated that the CD103^−^ and CD103^+^ CD4^+^CD69^+^ T cell populations can persist locally in the skin in the absence of continual replacement by circulating cells *(2),* thereby defining them functionally as T_RM_ populations.

Most human skin-resident memory T cells also express the cutaneous lymphocyte antigen (CLA), a glycan moiety that promotes skin homing of immune cells by acting as a ligand for E-selectin *(14).* CLA is also expressed by skin-tropic memory CD4^+^ T cells in human blood *(15),* but the clonal and functional relationship between the CD4^+^CLA^+^ T cells in blood and T_RM_ populations in the skin are not well defined. To elucidate the relationship between CD4^+^CLA^+^ T cells in the blood and skin, we characterized circulating and cutaneous T cell populations, taking advantage of new technological developments such as CyTOF, transcriptional profiling of small cell populations by RNA-sequencing, and novel humanized mouse models we have developed.

In tissue-explant and skin-humanized mouse systems, we showed that CD4^+^CLA^+^CD69^+^CD103^+^ T_RM_ in human skin can downregulate CD69 and exit the tissue. Furthermore, we identified a distinct population of CD4^+^CLA^+^CD69^−^CD103^+^ T cells in human lymph and blood that shared phenotypic, functional and transcriptomic signatures with the CD4^+^CLA^+^CD103^+^ T_RM_ population in the skin. TCR repertoire analysis confirmed the common clonal origins of the circulating and skin-resident CD4^+^CLA^+^CD103^+^ cells. In skin-humanized mice we further showed that following exit from the skin, CD4^+^CLA^+^CD103^+^ T cells could migrate into distant human skin sites and regain CD69 expression upon re-entering the tissue. Thus, basal recirculation of the CD4^+^CLA^+^CD103^+^ T_RM_ population can be detected in the steady state, and this can promote the spread of skin T_RM_ throughout this large barrier tissue.

## Results

### Cutaneous CD4^+^CLA^+^CD103^+^ T_RM_ cells can downregulate CD69 and exit the skin

Confirming prior analyses *(2)*, we found that the vast majority of both CD8^+^ and CD4^+^ T cells in human skin expressed CD69, and a subset of CD69^+^ cells also expressed CD103 and thus had the phenotype of cutaneous T_RM_ populations resistant to alemtuzumab-mediated depletion of circulating cells (Fig. 1, A and B; see Fig. S1 for representative T cell gating strategies from skin and blood). Consistent with their localization in the skin, most of these CD69^+^CD103^+^ cells also expressed CLA, (Fig. 1C) *(14)*. To directly assess the ability of different skin T cell populations to exit the tissue, we performed tissue explant cultures using human skin obtained from surgical samples. Despite high expression of CD69 in the tissue, we found that a fraction of cutaneous CD4^+^CLA^+^ T cells exited the tissue in these explant cultures and could be detected in the culture media. This tissue exit was associated with downregulation of CD69 by a fraction of both CD103^+^ and CD103^−^ cells (Fig. 1, D and E). Importantly, the input skin contained virtually no CD69^−^CD103^+^CD4^+^ T cells (Fig. 1D), and CD103 expression was not induced on blood-derived CD4^+^ T cells cultured in parallel (not shown), indicating that CD69 was indeed downregulated by CD103^+^ T_RM_ in these cultures. Additionally, the CD103^+^ cells found in the media did not express the chemokine receptor CCR7 (Fig. 1D), and thus these cells are distinct from CD69^−^CD103^+/lo^ cells that undergo CCR7-dependent migration from the skin to the draining lymph nodes in mice *(16),* and from the CCR7^+^CD62L^−^ migratory memory (T_MM_) cells described in human skin *(2).* By contrast, compared to CD4^+^ T cells, fewer CD8^+^ T cells exited the tissue in these skin explant cultures (Fig 1D), consistent with more prolonged tissue-residency of the skin CD8^+^ T_RM_ population in some mouse models *(7)(8)*.

**Figure 1:**
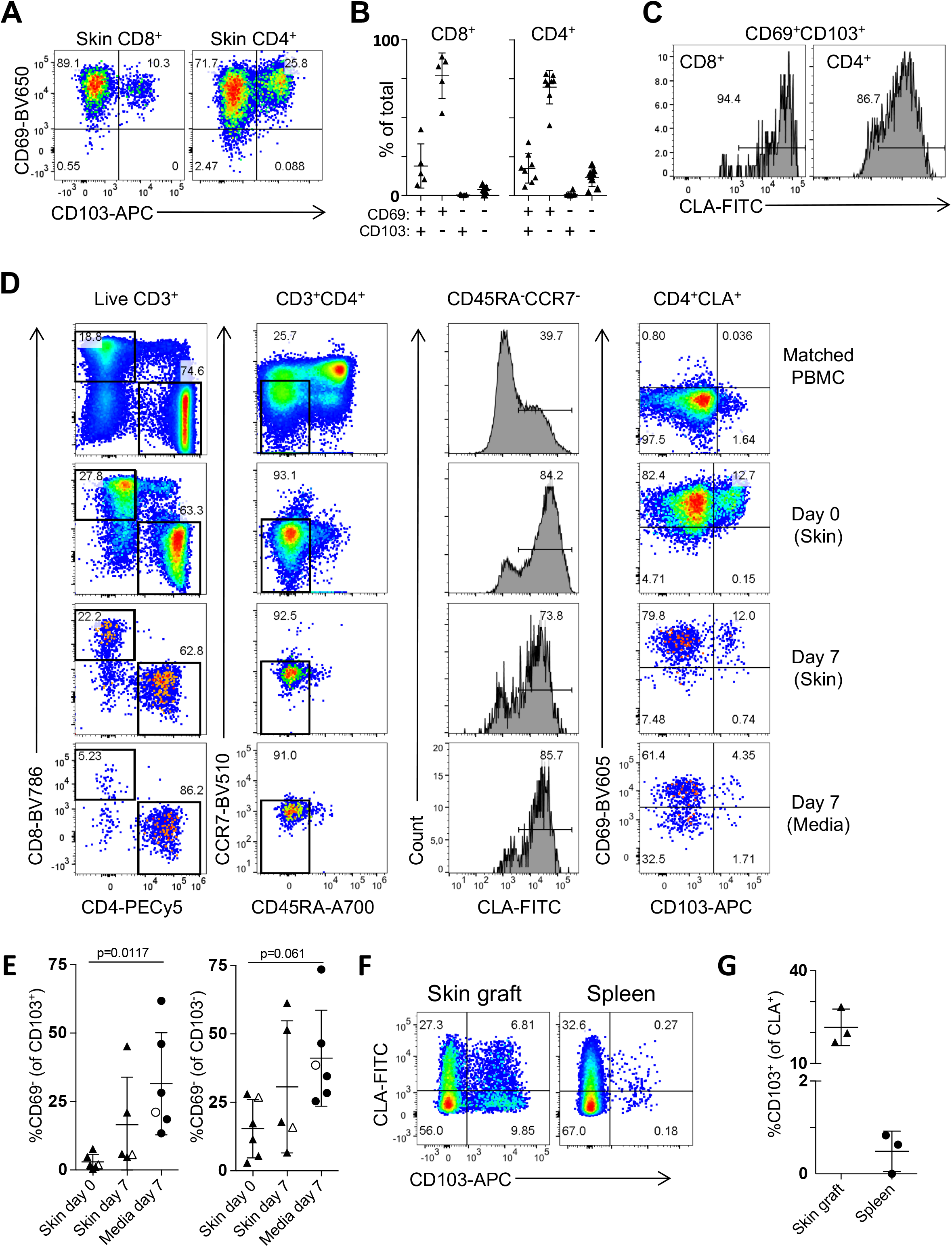
CD4^+^CLA^+^CD103^+^ T cells downregulate CD69 and exit the skin. **(A)** Representative flow cytometry analysis of CD69 and CD103 expression by live gated CD8^+^ and CD4^+^ T cells from human skin. **(B)** Graphical summary of the proportions of CD69- and CD103-defined T cell populations among CD8^+^ and CD4^+^ skin T cells. **(C)** Representative flow cytometry analysis of CLA expression by live gated CD103^+^CD69^+^ T_RM_ in human skin. **(D)** Human skin was adhered to tissue culture plates and cultured for 7 days submerged in media. The ratio of CD4^+^ and CD8^+^ T cells and the expression of CLA and CD103 by T cells in the indicated samples were analyzed by flow cytometry. Representative data (N=4). **(E)** Graphical summary of the proportion of CD69^−^ cells among CD103^+^ or CD103^−^ live gated CD45RA^−^ CD4^+^CLA^+^ T cells from the indicated samples. Open symbols represent data from a subject with mammary carcinoma but no skin condition. Significance determined by one-way repeated measures ANOVA with Tukey’s post-test for pairwise comparisons. **(F)** Three 8mm-punch biopsies of healthy human skin per animal (N=3) were placed on the back of NSG mice and grafts as well as spleens were analyzed by flow cytometry 50 days later. Representative flow cytometry analysis of CLA and CD103 expression by live gated human CD45^+^CD3^+^CD4^+^CD25^−^ CD45RA^−^ T cells. **(G)** Graphical summary showing CD103 expression by live gated human CD45^+^CD3^+^CD4^+^CD25^−^CD45RA^−^CLA^+^ T cells from skin grafts and spleens of skin-grafted NSG mice.

To determine if CD4^+^ T_RM_ could exit the skin *in vivo,* we used a xenografting model in which human skin was transplanted onto immunodeficient NOD, scid, common-*γ* chain-deficient (NSG) mice *(17).* T cells that had exited the skin were analyzed in the spleen. Similar to our explant studies, T cells exited the skin in all animals examined, including CLA^+^CD103^+^ T cells in 2 of the 3 recipient animals (Fig. 1, F and G). Importantly, expression of CD103 in the periphery is not induced by the xenogenic system we used, as we did not observe induction of CD103 expression by CD4^+^ T cells in NSG mice upon transfer of total PBMC (Fig. S2).

### Identification of CD4^+^CLA^+^CD103^+^ T cells in circulation of healthy humans

Among the CD4^+^CLA^+^CD45RA^−^ T cells in human blood, we noted a small population of CD103^+^CD69^−^ cells (Fig. 1D), and reasoned that these may represent CD4^+^CLA^+^CD103^+^ cells that had exited the skin and re-entered the circulation. To more comprehensively examine circulating CLA^+^CD103^+^ T cells and determine if they constitute a phenotypically distinct T cell population, we performed mass cytometry analysis of CLA^+^ T cells in the blood of 5 healthy subjects using markers associated with cell lineage, functional differentiation and migration of CD4^+^ T cells. Cluster analysis following t-distributed Stochastic Neighbor Embedding (t-SNE) revealed 10 clusters of CD3^+^CD45RA^−^CLA^+^ memory T cells present in all subjects examined (Fig. 2A, Fig. S3), including 5 clusters of CD4^+^ T cells (Fig. 2B). Interestingly, most CD4^+^CD103^+^ cells clustered together as a phenotypically discrete population (cluster 10). In addition to being positive for CD103 and its dimerization partner β7 integrin, cells in this cluster expressed chemokine receptors strongly indicative of skin tropism such as CCR4, CCR6, and CCR10, but were largely negative for CCR7 (Fig. 2, B and C) *(18)(19).* Additionally, cells in cluster 10 were low for expression of markers of regulatory T cells (Foxp3, CD25), T helper (Th)1 cells (CXCR3), Th17 cells (CD161) or natural killer T cells (CD56).

**Figure 2:**
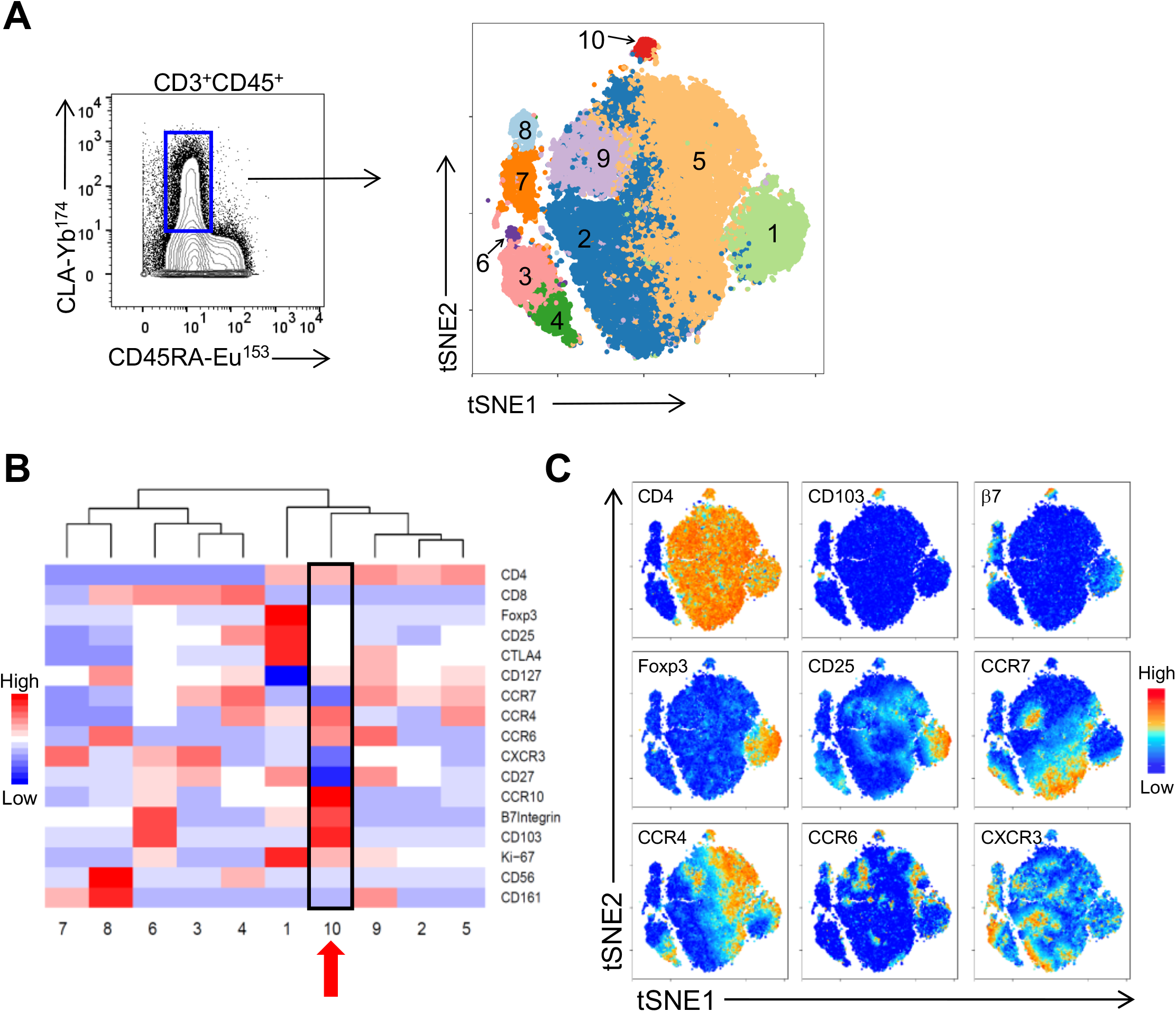
CD4^+^CLA^+^CD103^+^ T cells constitute a unique cell population in human blood. **(A)** (Left), Mass cytometry analysis of CD45RA and CLA expression by live gated CD3^+^CD45^+^ PBMC showing the gate used to define CD3^+^CLA^+^ T cells for subsequent clustering analysis. (Right), tSNE analysis and clustering of CD3^+^CLA^+^ T cells from blood of 5 healthy donors based on expression of CD4, CD8, CCR7, CD103, β7Integrin, CXCR3, CCR6, CCR4, CCR10, Foxp3, CD27, β7 integrin, CD25, CD161, and CD56. **(B)** Heat map showing relative expression of the indicated markers in each of the CD3^+^CLA^+^ cell clusters. **(C)** tSNE analysis of CD3^+^CLA^+^ T cells overlaid with relative expression of the indicated markers.

Using conventional flow cytometry, we directly compared the abundance and phenotype of circulating and skin-resident CD4^+^CLA^+^CD103^+^ T cells. In both blood and skin, CD103 and CCR7 clearly delineated 3 populations of CLA^+^ T cells (Fig. 3, A and B), but the distribution of these populations differed dramatically between sites. Whereas CD4^+^CLA^+^CD103^+^ memory T cells were common in the skin (26 +/− 9% of CD4^+^CLA^+^ T cells), they were a rare population in the blood, representing on average <2% of circulating CD4^+^CLA^+^CD45RA^−^ memory T cells, and <0.2% of total CD4^+^ T cells (Fig. 3C, Fig. S4A). Based on previous estimates of the total number of T cells in human skin and blood *(14),* we calculate that the number of CD4^+^CLA^+^CD103^+^ cells in the skin is approximately 250-fold higher than in the blood, highlighting the disequilibrium of this population between the skin and circulation (Fig. S4B). Within the skin, CD4^+^CLA^+^CD103^+^ cells were found in both the dermis and epidermis (Fig. S4C).

**Figure 3:**
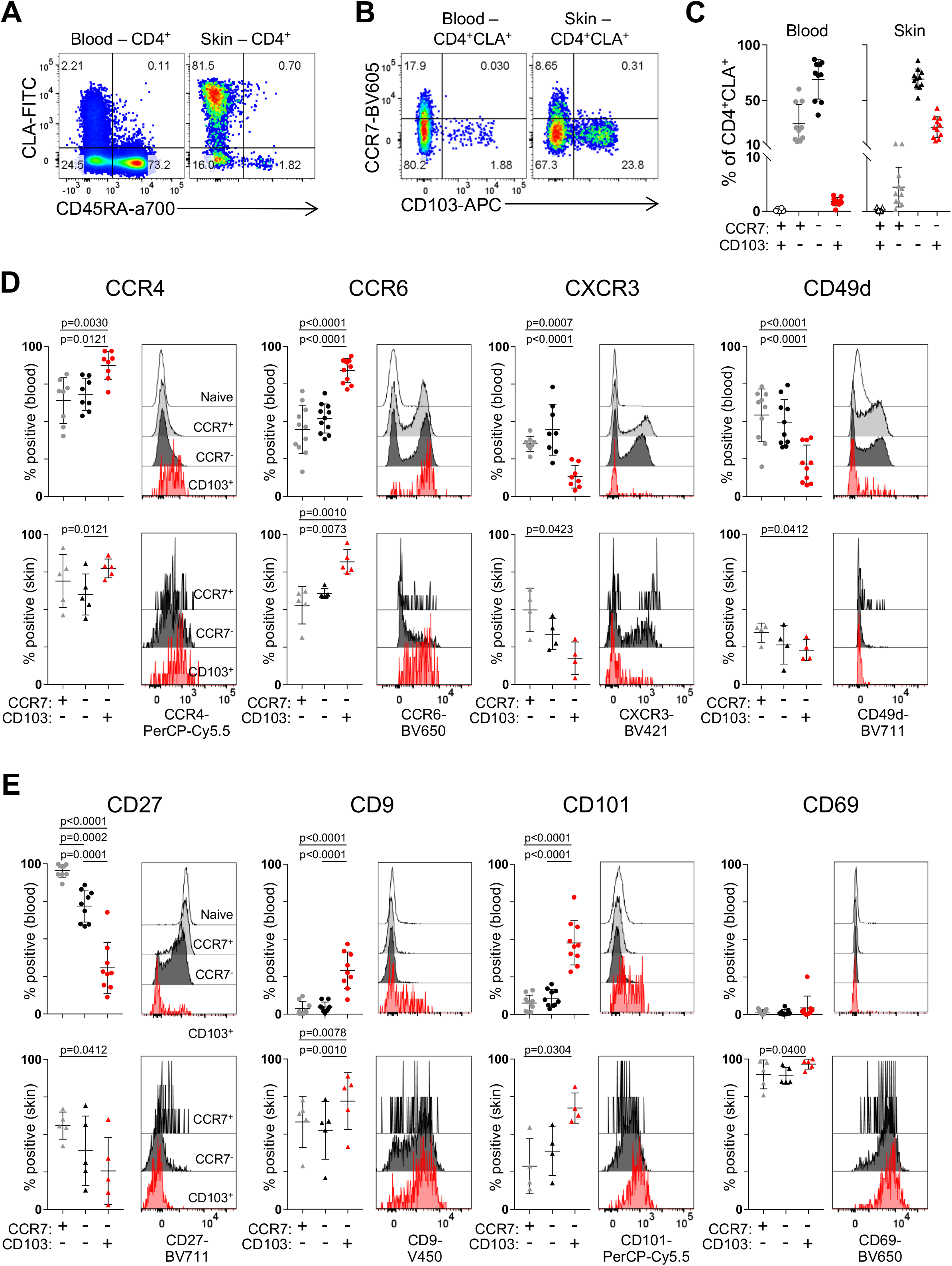
Shared phenotype of CD4^+^CLA^+^CD103^+^ T cells from human blood and skin. **(A)** Representative flow cytometry analysis of CD45RA and CLA expression by live gated CD4^+^ T cells from blood and skin of healthy donors. **(B)** Representative flow cytometry analysis of CCR7 and CD103 expression by live gated CD4^+^CD45RA^−^CLA^+^ memory T cells from blood and skin of healthy donors. **(C)** Graphical summary of the proportions of CCR7- and CD103-defined T cell populations among CD4^+^CD45RA^−^CLA^+^ cells from blood and skin. **(D,E)** Representative flow cytometry analysis and graphical summary of expression of the indicated markers by CD4^+^ T cell populations in the blood and skin as indicated. Significance determined by one-way repeated measures ANOVA with Tukey’s post-test for pairwise comparisons.

CD4^+^CLA^+^CD103^+^ T cells in the skin shared the CCR4^+^CCR6^+^CXCR3^−^ chemokine receptor profile with circulating CD4^+^CLA^+^CD103^+^ memory T cells. Additionally, CD4^+^CLA^+^CD103^+^ cells in the blood and all populations in the skin were largely negative for CD49d (Fig. 3D). Like CD103, CD49d (also known as α4 integrin) can pair with β7 integrin, and in this combination promotes T cell localization to the intestinal mucosa *(20).* CD4^+^CLA^+^CD103^+^ cells in the skin and blood were also CD27^−^ indicating that they are terminally differentiated *(21),* whereas most circulating CD4^+^CLA^+^CD103^−^ memory T cells were CD27^+^. Additionally, significant fractions of circulating CD4^+^CLA^+^CD103^+^ memory T cells expressed the markers CD101 and CD9, which were also expressed by the majority of CD4^+^CLA^+^CD103^+^ cells in the skin (Fig. 3E). Indeed, CD101 was recently identified as a marker of CD8^+^CD103^+^ T_RM_ populations in various human tissues that was also expressed by CD4^+^ lung T_RM_ *(3)(22),* whereas CD9 is highly expressed by keratinocytes and T cells in the skin, where it modulates TGF-β signaling, integrin function, cell migration, and wound healing *(23)(24)*. However, consistent with downregulation of CD69 by T cells that migrated out of the skin in our tissue explant cultures, CD4^+^CLA^+^CD103^+^ cells in the blood were almost all CD69^−^ (Fig. 3E).

### CD4^+^CLA^+^CD103^+^ T cells from blood and skin share a transcriptional and functional profile

To assess the transcriptional signature of circulating and skin-resident CD4^+^CLA^+^CD103^+^ T cells, we performed RNA-sequencing on sorted CLA^+^ memory CD4^+^ T cell populations from blood and skin (see Fig. S5 for sort scheme). Analysis of circulating CD4^+^CLA^+^CD103^+^ T cells identified a unique signature of 83 genes that were differentially expressed (false discovery rate (FDR) <0.05 and fold-change>2) compared to both CD4^+^CLA^+^CD103^−^CCR7^−^ and CD4^+^CLA^+^CD103^−^CCR7^+^ memory populations in the blood (Fig. 4A). This CD103^+^ gene signature derived from blood was also significantly enriched in CD4^+^CLA^+^CD103^+^ vs. CD4^+^CLA^+^CD103^−^CCR7^−^ T cells in the skin (Fig. 4B), consistent with the notion that circulating CD4^+^CLA^+^CD103^+^ cells represent skin T cells that downregulated CD69 to exit the tissue. Indeed, hierarchical clustering based on this gene signature grouped CD4^+^CLA^+^CD103^+^ cells from skin and blood into a single branch (Fig. 4C).

**Figure 4:**
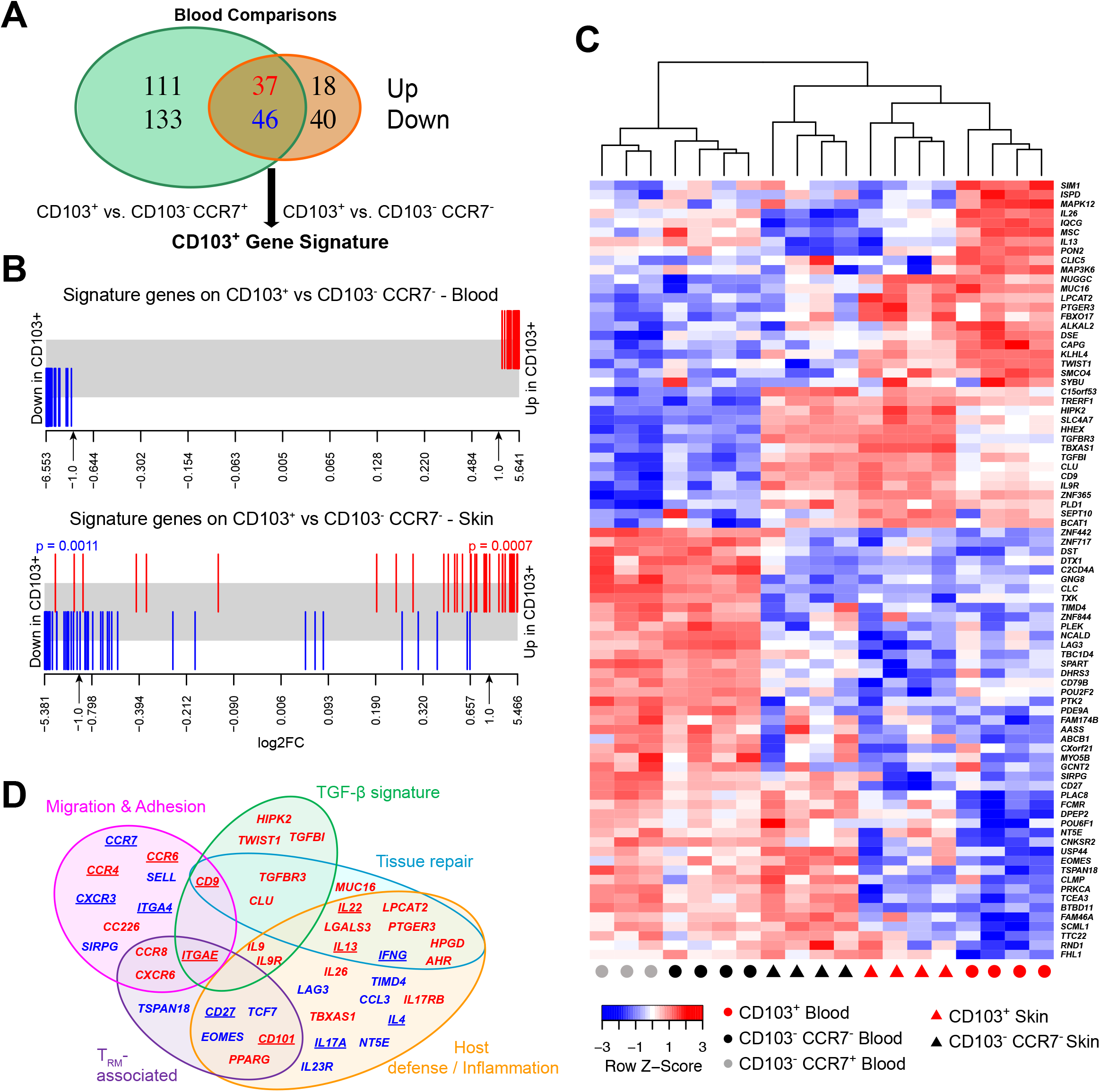
CD4^+^CLA^+^CD103^+^ T cells from human blood and skin share a transcriptional profile. Whole transcriptome profiling by RNA-sequencing was performed on sorted CLA^+^ T cell subsets from blood or skin. **(A)** Venn diagram showing the number of significantly differentially expressed (DE) genes (FDR<0.05 and log2 fold-change >1) between CLA^+^CD103^+^ T cells and either CLA^+^CD103^−^CCR7^+^ or CLA^+^CD103^−^CCR7^−^ T cells as indicated. The overlapping 83 genes were designated the CD103^+^ gene signature. **(B)** Barcode plot showing the distribution of the CD103^+^ signature genes (red=up-blue=down-regulated in CD103^+^) relative to gene expression changes comparing CD103^+^ and CD103^−^CCR7^−^ T cells from the blood or skin as indicated. Significance determined by rotation gene set testing for linear models. **(C)** Heat map and hierarchical clustering of RNA-seq samples from the indicated blood and skin cell populations based on the CD103^+^ gene signature. **(D)** Venn diagram showing functional annotation of key genes up- or down-regulated by CLA^+^CD103^+^ T cells in blood or skin identified in our phenotypic, functional, and transcriptional analyses. Category names were assigned based on described functions of the indicated genes in the published literature. Underlined gene names indicate proteins whose expression pattern was validated by flow cytometry in Figs 3 and 5.

CD103 expression by cutaneous T_RM_ is induced upon their migration into the skin by TGF-β *(25),* which is produced and activated by epidermal keratinocytes and dermal fibroblasts *(2)(26)(27).* Consistent with this, several TGF-β related genes were upregulated in both circulating and skin-resident CD4^+^CLA^+^CD103^+^ cells (Fig. 4D and Fig. S6). Moreover, along with *ITGAE* (the gene encoding CD103), *CD27* and *CD101*, we identified additional T_RM_-associated genes that were differentially expressed by circulating CD4^+^CLA^+^CD103^+^ cells, including *CCR8 (28), CXCR6 (3), EOMES (29) and PPARG (30).* We also identified overlapping functional modules of genes differentially expressed by CD4^+^CLA^+^CD103^+^ T cells that control cellular migration and adhesion, and that modulate host defense and tissue inflammation (Fig. 4D and Fig. S6).

To complement this transcriptomic study, we assessed the function of memory T cells from skin and blood following *ex vivo* stimulation and intracellular cytokine staining. Among effector cytokines, production of IL-22 and IL-13 was significantly enriched in CD4^+^CLA^+^CD103^+^ cells from both tissues, but these cells were largely negative for IFN-γ, IL-17A or IL-4 (Fig. 5). Although GM-CSF production was also highly enriched in CD4^+^CLA^+^CD103^+^ cells in the blood, production in the skin was low in all T cell populations (Fig. 5, D and E). This cytokine phenotype is consistent with that of Th22 cells *(31)(32)*, and distinguished CD4^+^CLA^+^CD103^+^ T cells from CD4^+^CLA^+^CD103^−^ and CD4^+^CLA^−^CD103^−^ T cells in both skin and blood. Co-production of IL-22 and IL-13 (Fig. S7) is indicative of a role for CD4^+^CLA^+^CD103^+^ T cells in promoting normal tissue homeostasis and repair in the skin *(33)(34)*. Consistent with this, CD4^+^CLA^+^CD103^+^ cells in the blood and skin differentially expressed a set of genes implicated in tissue-repair responses, such as *CD9 (24), MUC16 (35)(36),* and *LGALS3 (37)*, as well as the receptor components for the damage-associated epithelial cell products IL-25 *(IL17RA, IL17RB)* and prostaglandin E2 *(PTGER3)* (Fig. 4F, Fig. S6) *(38)(33)(39).* CD4^+^CLA^+^CD103^+^ T cells in both the skin and blood expressed *IL26,* a novel cytokine that can also have direct anti-microbial activity against *Staphylococcus aureus* and other extracellular bacteria *(40)(41).* Thus, the CD4^+^CLA^+^CD103^+^ T cell population is functionally well-suited to promote both tissue-repair and host-protective responses in the skin.

**Figure 5:**
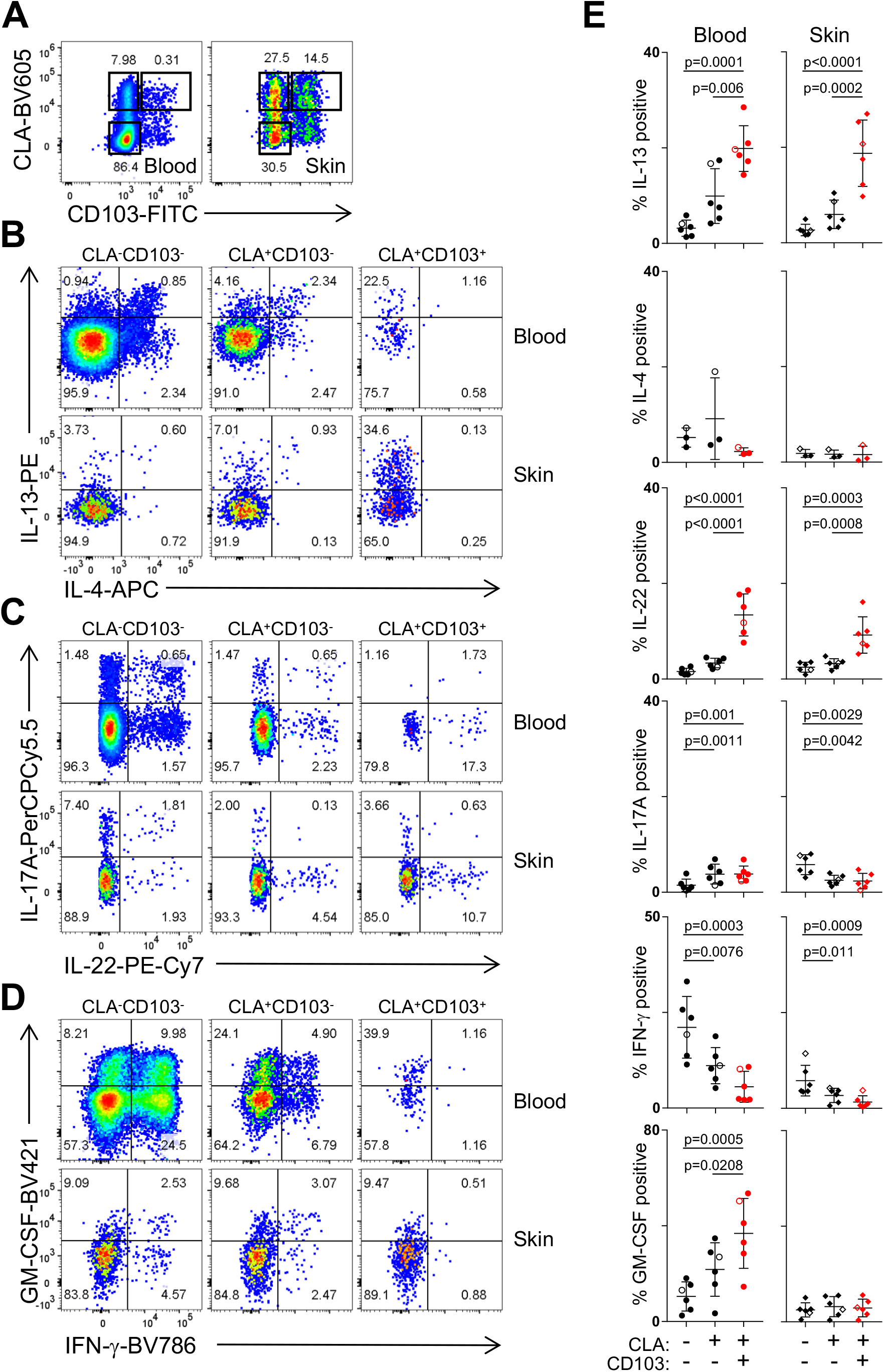
CD4^+^CLA^+^CD103^+^ T cells from human blood and skin share a functional profile. **(A)** Representative flow cytometry analysis of CD45RA and CLA expression by live gated CD4^+^ T cells from blood and skin of healthy donors. **(B,C,D)** Representative flow cytometry analysis of indicated CLA/CD103 subpopulations of blood and skin CD4^+^CD45RA^−^ cells producing IL-13, IL-4, IL-22, IL-17A, IFN-γ and GM-CSF as indicated upon *ex vivo* stimulation with PMA/ionomycin and intracellular cytokine staining. **(E)** Graphical summary of the proportions of CLA^−^CD103^−^, CLA^+^CD103^−^, and CLA^+^CD103^+^ CD4^+^CD45RA^−^ cells producing cytokines as indicated. Open symbols represent data from a subject with mammary carcinoma. Significance determined by one-way repeated measures ANOVA with Tukey’s post-test for pairwise comparisons.

### CLA^+^CD103^+^ T cells from human blood and skin are clonally related

Their shared phenotype, transcriptional signature, and functions suggest that the CD4^+^CLA^+^CD103^+^ T cells in blood and skin are closely related, and may represent circulating and skin-resident fractions of the same T_RM_ population. To directly determine if CD4^+^CLA^+^CD103^+^ T cells in the skin and blood have a shared clonal origin, we performed TCRβ sequencing on CD4^+^CLA^+^ memory CD4^+^ T cells from paired blood and skin samples from four individual donors that were sorted as in Fig. S5. Analysis of unique CDR3 clonotypes showed that sequences from CD4^+^CLA^+^CD103^+^ T cells from the skin were found at high frequency in the circulating CD4^+^CLA^+^CD103^+^ cells, and also showed some overlap with CD4^+^CLA^+^CD103^−^ skin T cells. By contrast, little clonal overlap was observed with circulating CLA^+^CD103^−^CCR7^+^ and CLA^+^CD103^−^CCR7^−^ T cells (Fig. 6A and Fig. S8A). Quantitative analysis of TCR repertoire overlap using the Morisita index *(42),* which accounts for both species presence and abundance, confirmed that the repertoire of skin CD4^+^CLA^+^CD103^+^ T cells is most similar to that of the circulating CD4^+^CLA^+^CD103^+^ T cell population (Fig 6A, right). Reciprocally, circulating CD4^+^CLA^+^CD103^+^ cells from the blood showed extensive TCR repertoire overlap with skin CLA^+^CD103^+^ T cells, but little clonal similarity with the other circulating CD4^+^CLA^+^ T cell populations (Fig 6B, Fig. S8A). Taken together, these analyses demonstrate the shared clonal origin of the CD4^+^CLA^+^CD103^+^ T cell populations in the blood and skin, thereby providing strong evidence that CLA^+^CD103^+^ T cells in the blood represent the circulating counterpart of the cutaneous CLA^+^CD103^+^ T_RM_ population. Notably, CD4^+^CLA^+^CD103^+^ T cells in both blood and skin showed diverse Vβ usage (Fig. S8B), and there was virtually no overlap in the TCRβ sequences in any of the populations examined between the 4 different donors (not shown). Thus, CD4^+^CLA^+^CD103^+^ T cells in the blood and skin do not appear to be a clonally restricted or invariant T cell population such as NK T cells or mucosa-associated invariant T (MAIT) cells *(43)*.

**Figure 6:**
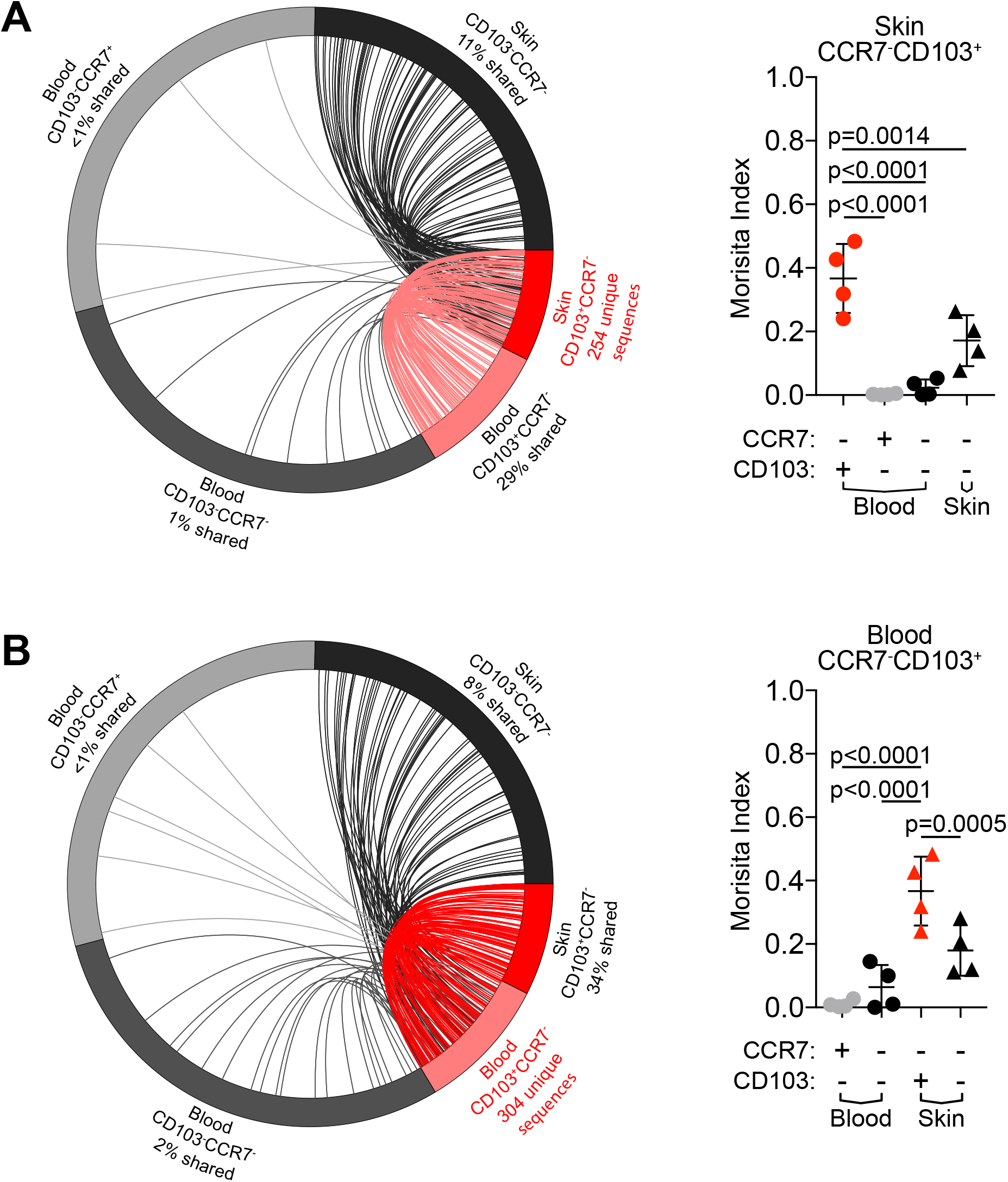
CD4^+^CLA^+^CD103^+^ T cells in skin and blood are clonally related. TCRβ-sequencing was performed on sorted CLA^+^ memory CD4^+^ T cell subsets (as in Fig. 4) from blood and skin sorted based on expression of CD103 and CCR7 as indicated. **(A)** (left) Circle plot of unique productive TCRβ sequences from each of the indicated populations of CLA^+^ T cells from one representative donor (donor 1). Connections highlight sequences from skin CLA^+^CD103^+^ cells found in each of the other populations. For visualization, sequences were downsampled (weighted for relative abundance) for populations containing >1000 unique sequences. (right) Graphical summary of the Morisita index as a measure of TCR repertoire similarity over all productive rearrangements between CLA^+^CD103^+^ cells in the blood and each of the indicated populations across all 4 donors examined. **(B)** Circle plot (left) and graphical summary of the Morisita index (right) as in **A** using CLA^+^CD103^+^ T cells from blood as the reference population. Significance determined by one-way ANOVA with Dunnett’s multiple comparisons test.

### CD4^+^CLA^+^CD103^+^ T cells recirculate from tissues via the lymphatics

To further determine if CD4^+^CLA^+^CD103^+^ cells recirculate from the tissue under physiological conditions, we analyzed human thoracic duct lymph (TDL) collected from chylothorax patients. As in the blood and skin, we identified a CD4^+^CLA^+^CD103^+^ population that lacked the expression of CCR7 (Fig 7A-C). Moreover, a majority of these cells expressed CCR4 and CCR6, but were low for CD27 and negative for CD69 expression (Fig. 7D). CCR7 is required for trafficking to the lymph node via the blood, and it is therefore unlikely that these CCR7^−^ T cells are recirculating by exiting the blood to the lymph node and then returning to the circulation via the TDL *(44).* Rather, this further supports the idea that circulating CD4^+^CLA^+^CD103^+^ T cells in blood and TDL have exited directly from the skin.

**Figure 7:**
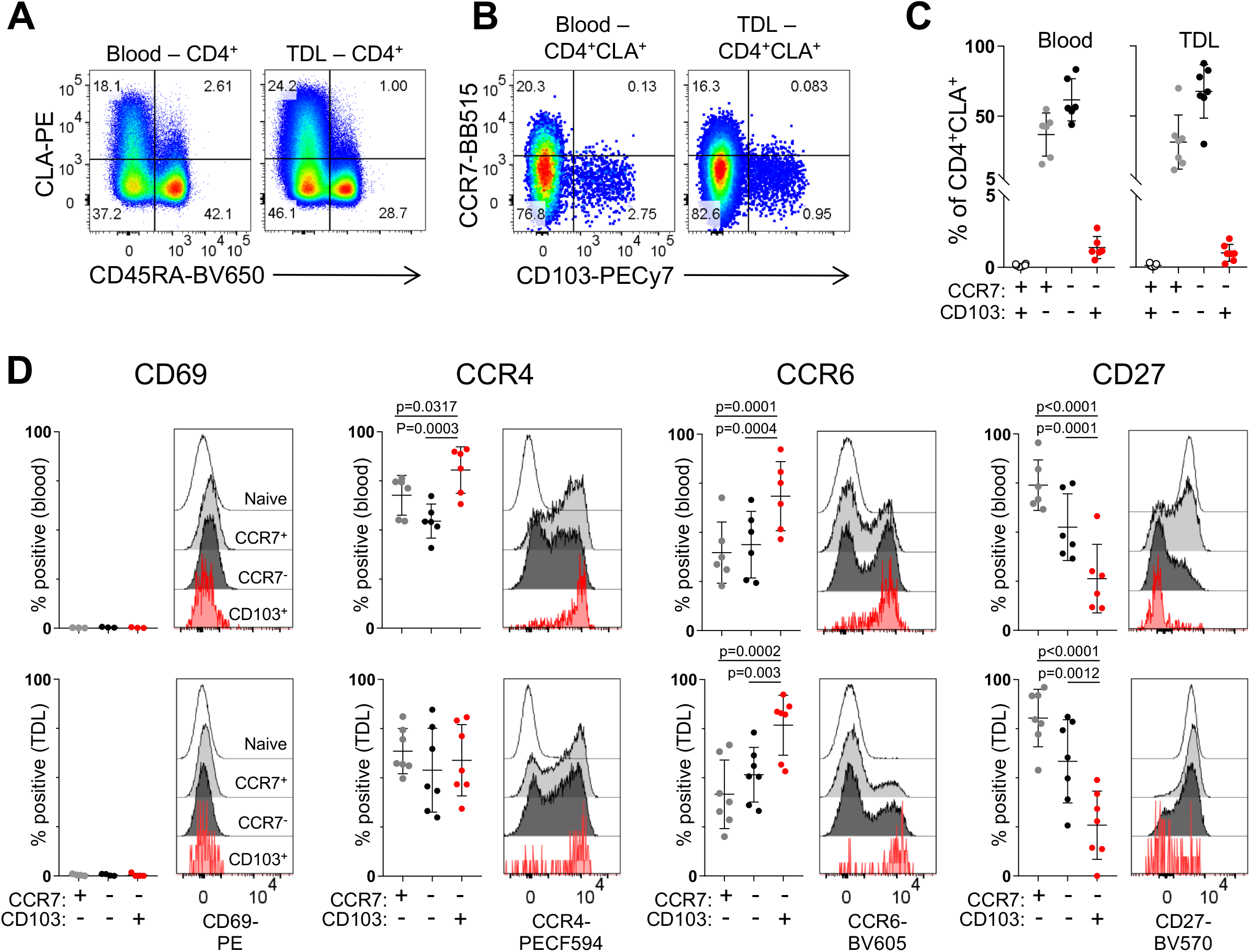
CD4^+^CLA^+^CD103^+^ T cells are present in human lymph. **(A)** Representative flow cytometry analysis of CD45RA and CLA expression by live gated CD4^+^ T cells from blood and TDL. **(B)** Representative flow cytometry analysis of CCR7 and CD103 expression by live gated CD4^+^CD45RA^−^CLA^+^ memory T cells from blood and TDL. **(C)** Graphical summary of the proportions of CCR7- and CD103-defined T cell populations among CD4^+^CD45RA^−^CLA^+^ cells from blood and TDL. **(D)** Representative flow cytometry analysis and graphical summary of expression of the indicated markers by CD4^+^ T cell populations in the blood and TDL as indicated. Significance determined by one-way repeated measures ANOVA with Tukey’s posttest for pairwise comparisons.

### Circulating CD4^+^CLA^+^CD103^+^ T_RM_ can reseed distant skin sites

Exit of cutaneous CD4^+^CLA^+^CD103^+^ T cells and their re-circulation may allow them to migrate to distant tissue sites, thereby promoting the efficient distribution of functionally specialized T cells throughout the skin. To directly test this hypothesis *in vivo*, we employed a mouse xenografting model designed to track tissue exit of human cutaneous T cells, and their subsequent migration to secondary human skin sites. In this system, cultured human keratinocytes and fibroblasts are placed in a grafting chamber that is surgically implanted on NSG mice. The cells undergo spontaneous cell sorting to form epidermal and dermal layers, generating engineered skin (ES) tissue with histological features of human skin as well as the organotypic expression of structural proteins such as human type VII collagen at the epidermal-dermal junction (Fig. 8A) *(45)*. Thus, the ES closely resembles human skin but lacks resident immune cells, and therefore T cell migration into the ES can be definitively monitored.

After healing of the ES (>110 days), mice received skin grafts from healthy donors, and tissues were analyzed three to five weeks later (Fig. 8, B and C). Similar to Figs 1F and 1G, CD4^+^CLA^+^CD103^+^ T cells had exited the skin grafts and were found in the spleens of all recipient animals. Additionally, in 5 of 7 recipient mice, CD4^+^CLA^+^CD103^+^ T cells were also found in the ES (Fig. 8, D and E), whereas no human cells were found in adjacent murine skin (Fig. S9). Similar to what we observed in human blood and skin, CD9 and CD69 were downmodulated on CD4^+^CLA^+^CD103^+^ T cells found in the spleen, but were re-expressed by cells entering the ES and CD27 expression remained low in all tissue sites (Fig. 8F). Finally, we used the ES system to interrogate the *in vivo* migration behavior of circulating CD4^+^CLA^+^CD103^+^ T cells from the blood. Upon transfer of PBMC into NSG mice carrying ES grafts (Fig. 8G), we found that T cells migrated to the ES, and that CD4^+^CLA^+^CD103^+^ cells were significantly enriched in the ES versus the spleen (Fig. 8, H and I). Together, these data demonstrate that CD4^+^CLA^+^CD103^+^ T cells that exit from the skin upon grafting or that are found in the blood of healthy individuals have the ability to migrate to and populate secondary skin sites.

**Figure 8:**
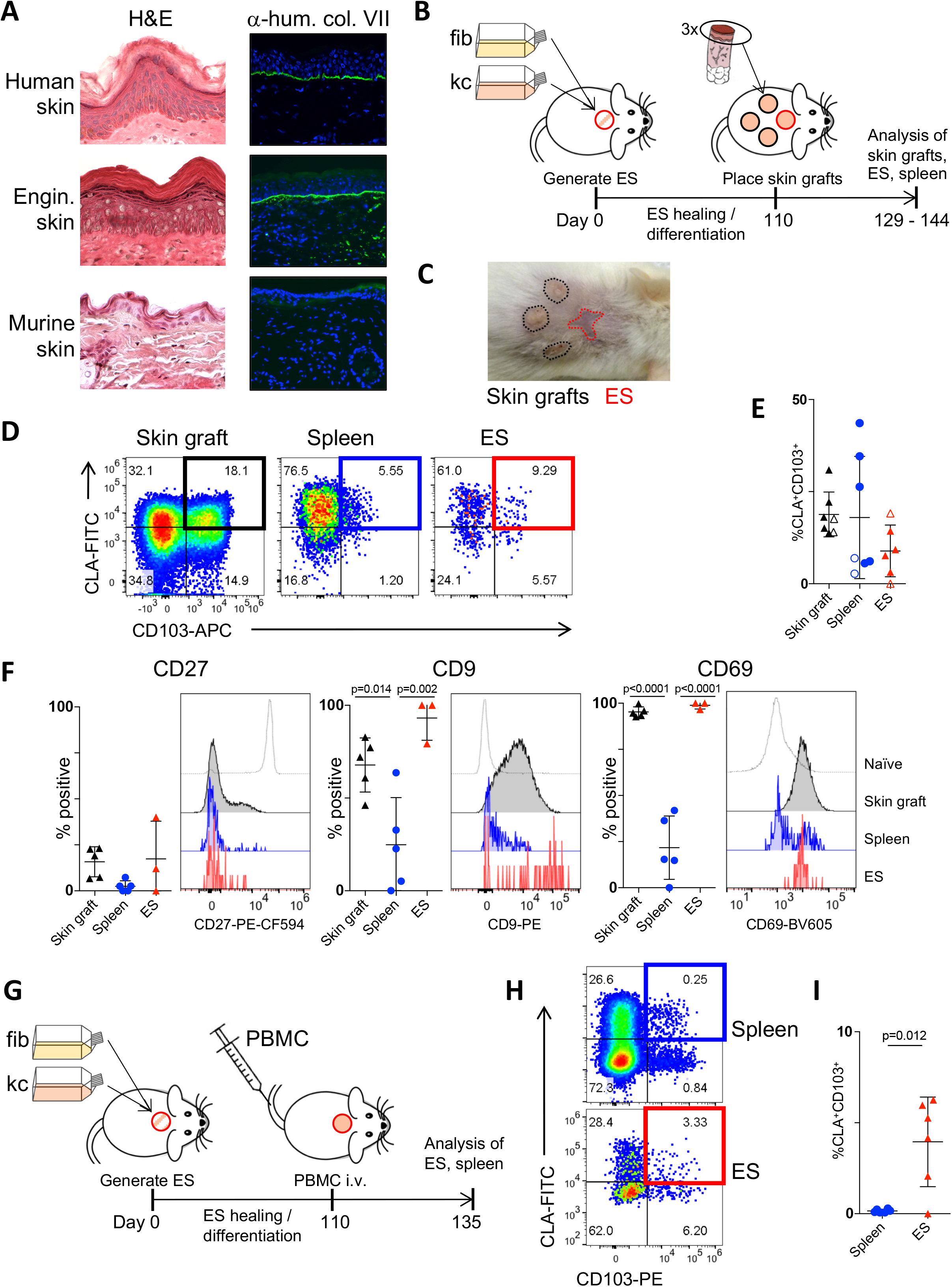
CD4^+^CLA^+^CD103^+^ T_RM_ can exit the skin and reseed distant skin sites in a xenograft model. **(A)** *In vitro* expanded human keratinocytes and fibroblasts were grafted onto the backs of NSG mice using a grafting chamber. After 99 days of healing and differentiation, the engineered skin or adjacent murine skin were excised, frozen in OCT and stained either with hematoxylin and eosin (left) or with anti-human type VII collagen prior to immunofluorescence analysis (right). Human skin from a healthy donor was used as control. **(B)** Experimental schematic for the generation of engineered skin (ES) followed by xenografting human skin onto NSG mice. **(C)** Representative photograph of ES and skin grafts on day 144. **(D)** Representative flow cytometry analysis and **(E)** graphical summary of CLA^+^CD103^+^ cells by live gated human CD45^+^CD3^+^CD4^+^CD45RA^−^ T cells from skin grafts, spleen, and ES (3-5 weeks after skin grafting). Open and filled symbols denote samples derived from 2 different skin donors. Each symbol represents data from one recipient animal. **(F)** Representative flow cytometry analysis and graphical summary of expression of CD27, CD9 and CD69 by live gated CD45^+^CD4^+^CD45RA^−^CD103^+^CLA^+^ T cells in the skin grafts, spleen, and ES 5 weeks after skin grafting (day 145 relative to ES generation). Significance determined by one-way ANOVA with Tukey’s post-test for pairwise comparisons. **(G)** Experimental schematic for the generation of engineered skin (ES) followed by adoptive transfer of 2.5×10^6^ PBMC (autologous to the ES)/mouse into NSG mice. (H) Representative flow cytometry analysis and (I) graphical summary of CLA^+^CD103^+^ cells by live gated human CD45^+^CD3^+^CD4^+^CD45RA^−^ T cells from spleen, and ES 25 days after PBMC transfer. Each symbol represents data from one recipient animal. Significance determined by paired t-test.

## Discussion

T_RM_ populations mediate optimal protective responses to site-specific challenges in nonlymphoid tissues, and are most readily identified by their expression of CD69 and/or CD103 *(4)(2).* Using tissue explant cultures and a skin-xenografting mouse model, we made the surprising discovery that CD4^+^CLA^+^CD69^+^CD103^+^ T cells in human skin can downregulate CD69 and exit the tissue. This is consistent with recent observations in murine systems showing that secondary stimulation mobilized CD8^+^ T cells from non-lymphoid tissues (including the skin) that subsequently established residence within draining secondary lymphoid organs (SLO) *(46).* However, this study did not establish from which skin-resident population these cells were derived, or whether these mobilized cells could further recirculate to other peripheral nonlymphoid tissue sites. We also identified CD4^+^CLA^+^CD103^+^ T cells as a distinct population of circulating T cells that are clonally related to the CD4^+^CLA^+^CD103^+^ T_RM_ population in the skin. Importantly, the phenotypic, functional and transcriptional profile of circulating CD4^+^CLA^+^CD103^+^ T cells is consistent with their origin and function within the skin, a TGF-β-rich barrier site exposed to microbial threats and frequent tissue damage. Finally, we show that upon exiting the skin, CLA^+^CLA^+^CD103^+^ T cells can migrate via the circulation to secondary skin sites where they re-acquire markers of tissue residency such as CD69. Based on these features, we propose that blood and skin CD4^+^CLA^+^CD103^+^ T cells represent components of the same T_RM_ population that undergoes basal recirculation, but is maintained in substantial disequilibrium between these tissues.

Our data challenge current concepts regarding the strict tissue compartmentalization of T cell memory in humans, and instead support a model in which cells of the CD4^+^CLA^+^CD103^+^ T_RM_ population can transiently forgo their tissue-location to re-assume residency at distant sites of the tissue. Whether tissue-exit of cutaneous CD4^+^CLA^+^CD103^+^ T cells is a stochastic process or actively triggered mobilization remains to be determined. In the context of our studies in explant cultures and in skin-humanized mice, tissue damage unavoidably associated with surgical skin acquisition is one potential trigger that may have impacted T_RM_ mobilization. However, we detected CD4^+^CLA^+^CD103^+^ cells in the blood and lymph of all donors analyzed, which indicates that a small fraction of the CD4^+^CLA^+^CD103^+^ T_RM_ population recirculates even in the absence of clinical skin infection, inflammation or tissue damage.

Our xenografting studies allowed us to further follow the fate of tissue-derived T cells, and we found that CD4^+^CLA^+^CD103^+^ T cells from either skin grafts or from human blood can migrate to and preferentially seed secondary human skin sites. This tissue-seeding occurred in the absence of tissue damage or local inflammation since the recipient engineered skin tissue was fully healed (>110 days) and thus lacked expression of damage-associated molecules such as IL-1α, IL-1β, IL-18 and TNF-α that might increase recruitment of circulating cells *(47).* However, it remains possible that recruitment to secondary skin sites is increased by damage-associated stress. This might offer an intriguing explanation to the hitherto unexplained Koebner phenomenon, in which lesions in T_RM_-mediated diseases such as psoriasis and mycosis fungoides can spread to otherwise healthy (non-infected) skin sites upon triggers such as mechanical trauma, burns, friction or UV-irradiation *(48)(49)(50).*

Human skin-resident αβ T cells can promote tissue-repair in skin organ culture models *(51).* Although the precise role of the CD4^+^CLA^+^CD103^+^ population in skin immunity and homeostasis remains to be established *in vivo,* their transcriptional and functional profile is indicative of a function in wound-healing and tissue-repair responses. The signature cytokines produced by CD4^+^CLA^+^CD103^+^ T cells, IL-22 and IL-13, both have important tissue-repair functions in the skin. IL-22 acts directly on keratinocytes to promote their survival, proliferation, migration and anti-microbial functions *(34),* whereas IL-13 activates cutaneous fibroblasts and promotes M2 macrophage differentiation and wound healing *(52).* IL-22 can also induce production of anti-microbial peptides by keratinocytes *(53).* Thus, the CD4^+^CLA^+^CD103^+^ population has many of the hallmarks of other lymphocyte populations implicated in antimicrobial and tissue-repair responses, such as cutaneous IL-22 producing γδ T cells, and IL-13- or IL-22-producing ILC cells *(54)(55)(34).* In these contexts, it is important to note that we and others found CD4^+^CLA^+^CD103^+^ T cells in the epidermis and the dermis of the skin (2), and thus they are ideally positioned to modulate the responses of keratinocytes, fibroblasts and skin macrophages and promote both tissue-repair and host-protective responses.

Mobilization of cutaneous CD4^+^CLA^+^CD103^+^ T cells to the circulation would support the distribution of immunity in a large barrier organ such as the skin, as well as provide a reservoir of specialized circulating T cells that could be rapidly recruited to infected or damaged skin to promote host-defense and tissue-repair. Importantly, our identification of CD4^+^CLA^+^CD103^+^ T cells as a unique population of circulating T cells in healthy subjects greatly facilitates the isolation and study of cutaneous T_RM_ from a broadly available human tissue, the blood. This promises to yield new insights into the biology and function of the human skin T_RM_ population, and opens novel avenues to therapeutically manipulating skin T_RM_ in the contexts of cutaneous autoimmunity, infection, and tissue-repair.

## Material and Methods

### Study design

The objective of this research was to characterize the phenotype, function and migratory behavior of CD4^+^ T cell populations that express the cutaneous lymphocyte antigen (CLA), and to define the relationship between CLA^+^ T cells in the blood and CLA^+^ T_RM_ cells in the skin. This was accomplished using blood and skin samples from healthy donors by flow cytometric analysis of cellular phenotype and function, transcriptomic analysis by RNA-sequencing, T cell receptor clonotype analysis by TCR-sequencing, and experimental studies of cellular behaviour in explant culture models and skin xenograft studies using immunodeficient mice. Blood samples from healthy donors was obtained by standard phlebotomy. Normal human skin was obtained from patients undergoing elective surgery (panniculectomy, elective breast reduction), in which skin was discarded as a routine procedure. In one case, skin and blood was obtained from a treatment naïve subject undergoing surgery for mammary carcinoma, and data from this subject is specifically marked in the figures. Samples of subjects of both sexes were included in the study. Ages ranged from 17 – 70. All samples were obtained upon written informed consent at the University Hospital Salzburg, Austria, the University of Pennsylvania and the Children’s Hospital of Philadelphia (Philadelphia, PA) or, or the Virginia Mason Medical Center in Seattle, WA, USA. All studies were approved by the Salzburg state Ethics Commission (decision: according to Salzburg state hospital law no approval required) (Salzburg, Austria), or the Institutional Review Board of the University of Pennsylvania and the Children’s Hospital of Philadelphia (Philadelphia, PA), or the Institutional Review Board of the Benaroya Research Institute (Seattle, WA). NOD.Cg-Prkdcscid Il2rgtm1Wjl/SzJ (NSG) mice were obtained from The Jackson Laboratory and bred and maintained in a specific pathogen-free facility in accordance with the guidelines of the Central Animal Facility of the University of Salzburg. All animal studies were approved by the Austrian Federal Ministry of Science, Research and Economy. No statistical method was used to predetermine sample size. Samples sizes were based on the availability of human blood and skin specimens and were large enough to ensure achieve a greater than 80% probability of identifying an effect of >20% in measured variables. 2-3 independent experiments (biological replicates) were conducted to validate each finding. In RNA sequencing, samples were excluded from the analysis based on pre-established quality control criteria: Samples with a total number of fastq reads <1×10^6^, mapped reads below 70% or median CV coverage >1 were excluded from further analysis. Randomization was not applicable because no treatment or intervention groups were included in the study. Blinding was not applicable since no treatment groups were compared.

### Skin explant cultures

Skin was washed in PBS with 1% Pen/Strep and 0.1% Primocin (Invivogen; ant-pm-1) for 5 minutes. Small skin pieces of 1-2mm were generated using forceps and sharp scissors. Pieces were placed in 60mm dish and allowed to adhere for 10 minutes. Crawl-out medium consisting of 50% Epilife (Gibco, MEPICF500) and 50% RPMI-complete (RPMIc: RPMI 1640 (Gibco; 31870074) with 5% human serum (Sigma-Aldrich; H5667 or H4522), 1% penicillin/streptomycin (Sigma-Aldrich; P0781), 1% L-Glutamine (Gibco; A2916801), 1% NEAA (Gibco; 11140035), 1% Sodium-Pyruvate (Sigma-Aldrich; S8636) and 0.1% β-Mercaptoethanol (Gibco; 31350-010)) was added to explant cultures. Seven days later, cells in culture medium were analyzed by flow cytometry.

### Cytometry by time-of-flight (CyTOF)

Human frozen PBMCs were thawed and rested for 12-15h. The samples were washed with Ca- and Mg-free PBS (Sigma; D8537) and stained with 50μM Cisplatin (Enzo Life Sciences; ALX-400-040-M250) in PBS for 1 minute to exclude dead cells. The cells were washed and resuspended with Human TruStain FcX (Biolegend; 422302) for five minutes before adding the primary surface staining cocktail for 20m, washing and staining with the secondary surface cocktail for 20m. Intracellular staining was performed following fixation and permeabilization using the Maxpar Nuclear Antigen Staining Buffer Set (Fluidigm; 201063) for 60 minutes, after which cells were incubated overnight at 4°C with Maxpar Fix and Perm Solution containing 125nM Cell-ID Intercalator-Ir (Fluidigm; 201192A) for DNA staining. Cells were washed with MilliQ H_2_O and resuspended in MilliQ H_2_O spiked with 1/20th Maxpar EQ Four Element Calibration Beads (Fluidigm; 201078) to a density of <500.000 cells/mL. Data was acquired on a CyTOF 1.5 (Fluidigm) instrument. For analysis, FCS files on gated CD3^+^CLA^+^ cells were generated using FlowJo software (Tree Star, Inc.). t-distributed Stochastic Neighbor Embedding (t-SNE) analysis was performed in R using the cytofkit R package, and clustering was performed with FlowSOM.

### T cell isolation from skin for flow cytometry and RNAseq

Tissues were minced and digested using collagenase type 4 and DNase as previously described *(56)*. Briefly, subcutaneous fat was removed before skin was minced with dissection scissors and surgical scalpel. Approximately 1cm^2^ of skin was digested overnight in 5%CO2 at 37°C with 3ml of digestion mix containing 0.8mg/ml Collagenase Type 4 (Worthington; #LS004186) and 0.02mg/ml DNase (Sigma-Aldrich; DN25) in RPMIc. Samples were filtered, washed with RPMI and PBS and stained for flow cytometry/cell sorting or stimulated for intracellular cytokine staining. Cryopreserved PBMC or TDL cells were rested overnight at 37°C and 5% CO2 in complete medium (RPMI supplemented with 10% FBS, 2 mM L-glutamine, 100 U/ml penicillin, and 100 mg/ml streptomycin), washed and stained for flow cytometry.

### PBMC isolation for flow cytometry and RNAseq

Human PBMCs were isolated using Ficoll-Hypaque (GE-Healthcare; GE17-1440-02) gradient separation. For RNAseq CD4^+^ T cells were enriched using CD4 microbeads (Miltenyi; 130-045-101) and 4×10^6^ cells/ml were resuspended in RPMIc with 50 U/ml IL-2 (Immunotools; 11340023) in a 24-well and stimulated for 20h with 25μl/ml ImmunoCult CD3/CD28 T cell activator (Stemcell; 10971) prior to staining and cell sorting.

### Flow cytometry

For sorting, cells were stained in FACS buffer (PBS + 1 % FBS +1 mM EDTA) for surface markers. For detection of intracellular cytokine production, skin single cell suspensions and PBMCs were stimulated with 50 ng/ml PMA (Sigma-Aldrich; P8139) and 1 μg/ml Ionomycin (Sigma-Aldrich; I06434) with 10 μg/ml Brefeldin A (Sigma-Aldrich; B6542) for 3.5 hrs. For permeabilization and fixation Cytofix/Cytoperm kit was used (BectonDickinson; RUO 554714). Data were acquired on LSR Fortessa, LSRII, FACSymphony (all BD Biosciences) or Cytoflex LS (Beckman Coulter) flow cytometers and analyzed using FlowJo software (Tree Star, Inc.).

### Analysis of thoracic duct lymph (TDL) samples

Thoracic duct lymph was collected and processed from patients with chylopericardium and chylothorax as previously published *(57)(58)*. Cells were washed in PBS and stained for CXCR3, CCR4, CCR6, and CCR7 at 37°C, 5% CO2 for 20 minutes. All subsequent incubations were performed in the dark at room temperature. Cells were stained for viability exclusion for 10 minutes. Antibody cocktail mix was added and incubated for 20 minutes. Cells were fixed with PBS with 1% paraformaldehyde and stored in the dark at 4°C overnight until acquisition. Data were collected on a BD FACSymphony flow cytometer (BD Biosciences) and analyzed using FlowJo software (Treestar).

### RNA sequencing

500 cells per population were sorted into lysis buffer using a BD FACSARIA III or FACSFUSION instruments (BD Biosciences), and cDNA was prepared using the SMART-Seq v4 Ultra Low Input RNA Kit for Sequencing (Takara). Library construction was performed using the NexteraXT DNA sample preparation kit (Illumina) using half the recommended volumes and reagents. Dual-index, single-read sequencing of pooled libraries was run on a HiSeq2500 sequencer (Illumina) with 58-base reads and a target depth of 5 million reads per sample. Base-calling and demultiplexing were performed automatically on BaseSpace (Illumina) to generate FASTQ files.

### RNAseq analysis

The FASTQ files were processed in order to remove reads of zero length (fastq_trimmer v.1.0.0), remove adapter sequences (fastqmcf tool v.1.1.2) and perform quality trimming from both ends until a minimum base quality ≥ 30 (FASTQ quality trimmer tool v.1.0.0). Reads were aligned to the human reference genome (build hg38) with TopHat (v.1.4.0) and read counts per Ensembl gene ID were quantified with htseq-count (v.0.4.1). Quality metrics for the FASTQ and BAM/SAM files were generated with FastQC (v.0.11.3) and Picard (v.1.128). Processing of FASTQ and BAM/SAM files was executed on the Galaxy workflow platform of Globus genomics. Statistical analysis of gene expression was assessed in the R environment (v.3.4.4). Samples with a total number of fastq reads below 10^6^, mapped reads below 70% or median CV coverage > 1 were excluded from further analysis. Mapping Ensembl Gene IDs to HGNC gene symbols was achieved through biomaRt (GRCh38.p10). Genes were filtered for protein coding genes and those with an expression of CPM > 2 in at least 10% of the libraries. A linear model for gene expression was fit to the filtered 12,293 genes using limma (v3.34.9)(59), considering donor effects through a random factor. For visualizations the random effect of the model was approximated by removing the donor effect from the expression data with limma::removeBatchEffect. Genes found to be significantly different (adj.p.val < 0.05 and fold-change>2) between CD103^+^ cells and CD103^−^CCR7^+^ cells as well as between CD103^+^ cells and CD103^−^CCR7^−^ cells in the blood were defined as the CD103^+^ gene signature. Enrichment of the CD103^+^ gene signature in the ranked list of CD103^+^ cells vs CD103^−^CCR7^−^ cells in the skin was visualized with limma::barcodeplot and significance determined by rotation gene set testing with limma::roast. Publicly available data on human CD4^+^ Tissue-Resident Memory T Cells was retrieved from GEO dataset GSE94964. Gene names were lifted over from hg19 to hg38 nomenclature, the limma::voom transformed expression values were corrected for donor effect with sva::ComBat and PCA was performed on genes from the CD103^+^ signature or the same number of randomly selected genes.

### TCRβ sequencing and analysis

A minimum of 2,000 cells T cells from the indicated populations were sorted into RPMIc, and genomic DNA was prepared using the QIAamp DNA Micro Kit (Qiagen). Amplification and sequencing was performed using the immunoSEQ^®^ Assay (Adaptive Biotechnologies, Seattle, WA), which combines multiplex PCR with high throughput sequencing and a bioinformatics pipeline for TCRβ CDR3 analysis *(60)*. Data analysis was performed using Adaptive Biotechnologies ImmunoSeq Analyzer 3.0 software and R version 3.5.1. Data for the analysis in R was exported through the export function in the Rearrangement details view. Circle plots for individual donors were created by downsampling populations with more than 1000 unique rearrangements (weighted based on relative abundance of each individual clonotype) and matching TCR chains with the R package TCRtools (https://github.com/mjdufort/TCRtools). Links between the blood or skin CD103^+^ reference population and all other populations are displayed in the circle plot. For V gene usage, we removed unknown and ambiguous mappings and computed the percentage of clones using each V gene among each sample. The plot includes V genes that have a usage ≤ 5% in at least one sample.

### Generation of engineered skin (ES)

Human keratinocytes and fibroblasts were isolated from normal human skin and immortalized using human papilloma viral oncogenes E6/E7 HPV as previously described *(61)*. These were cultured in Epilife (Gibco, MEPICF500) and DMEM (Gibco; 11960-044) containing 2% L-Glutamine, 1% Pen/Strep, 10% FBS, respectively. For transplantation, 80% confluent cells were trypsinized (TrypLE express, Gibco; 12604021) washed with PBS and counted. Per mouse, 1-2×10^6^ keratinocytes were mixed 1:1 with autologous fibroblasts in 400μl MEM(Gibco; 11380037) containing 1% FBS, 1% L-Glutamine and 1% NEAA were used for *in vivo* generation of engineered skin as described *(45).*

### Transplantation of human skin or PBMC transfer

8mm punch biopsies of human skin were trimmed to an average thickness of 0.5-1mm. Transplants were soaked in PBS + 1% Pen/Strep + 0. 1% Primocin (Invitrogen; ant-pm-1) for 5 minutes and kept on ice in a sterile container with PBS soaked gauze until transplantation. NSG mice were anesthetized and full thickness wound bed prepared using surgical scissors. Three grafts/mouse were placed on the back and fixed using surgical skin glue (Histoacryl; BRAUN). Transplants were covered with paraffin gauze dressing and bandaged with self-adhesive wound dressing. Bandages were removed after 7 days. In other experiments PBMC were thawed and rested in media over night before transfer into NSG recipient mice (2.5×10^6^ cells/animal). After transfer of human cells (PBMC or skin grafting) mouse neutrophils were depleted with anti-Gr-1 antibody (InVivoMab clone RB6-8C5; 100 μg Gr-1-Ab/animal i.p. every 5-7 days) *(62)(63).*

### Histological staining of skin sections

Normal human skin, engineered skin grafts and adjacent murine skin was excised and frozen in TissueTek O.C.T. Compound (Sakura; TTEK). 7 μm cryosections were stained with Hemalum solution acid (Carl Rorth; T865.1) and Eosin Y aqueous solution (Sigma, 201192A). Human type VII collagen was stained by immunofluorescence using anti-human type VII collagen antibody (anti-NC-1 domain of type VII collagen (LH7.2) kindly provided by Dr. Alexander Nyström, University of Freiburg, Germany) and goat anti-rabbit A488 (ThermoFisher; A11008) secondary antibody, and nuclear DAPI staining (ProLong™ Gold Antifade Mountant with DAPI, Invitrogen; P36931).

### Tissue preparation from mice

Mice were euthanized using CO2 asphyxiation followed by cervical dislocation. Single cell suspensions were generated from spleen, engineered skin and human skin grafts and leukocytes analyzed by flow cytometry.

### Statistical analysis

Statistical significance of data was calculated with Prism 6.0 software (GraphPad) by one-way ANOVA with Tukey’s or Dunnett’s multiple comparisons test, or by paired t-test as indicated. Error bars indicate mean + standard deviation.

## Supplementary Materials

### Supplementary Data Methods

#### T cell isolation from murine skin for flow cytometry

Approx. 3 cm^2^ of shaved dorsal mouse skin were harvested and single cell suspensions prepared as in *(64)* and stained for flow cytometry.

#### Separation of epidermis and dermis

Skin was washed in PBS with 1% Pen/Strep and 0.1% Primocin for 5 minutes prior to removal of fat tissue. Skin pieces of 2-3 mm in diameter were incubated overnight with 0.25% Dispase II (Sigma-Aldrich; D4693) in RPMI-complete and dermis and epidermis separated with forceps. The separated tissue layers were then further minced and digested with collagenase type 4 and DNase to generate single cell suspensions for flow cytometric analysis.

### List of Supplementary Figures

Fig. S1: Representative flow cytometry gating strategies used to identify T cell subsets in human blood and skin.

Fig. S2: CD103 expression is not induced on human CD4^+^ T cells in NSG mice.

Fig. S3: CyTOF analysis of CLA^+^ T cells in PBMC.

Fig. S4: Frequencies of CLA^+^ T cell subsets in blood and skin.

Fig. S5: Experimental schematic of cell isolation and sort gates for RNA-seq.

Fig. S6: Graphical analyses showing expression of the indicated genes in the different populations of CD4^+^CLA^+^ T cells from blood and skin as determined by RNA-seq.

Fig. S7: CD103^+^CLA^+^ T cells from human blood and skin coproduce IL-22 and IL-13.

Fig. S8: Analysis of TCRβ repertoire overlap and Vβ gene usage in CLA^+^ T cells from blood and skin.

Fig. S9: Human skin-derived cells do not infiltrate murine skin.

Table S1: Detailed list of antibodies and reagents

## Acknowledgements

We thank Dr. A. Sir and Dr. R. Reitsamer of the Breast Center of the University Hospital Salzburg, Paracelsus Medical University Salzburg, Austria for providing us with skin and blood samples. We also thank Anna Hochreiter, MSc, and Anna Raninger, MSc, from the Flow Cytometry Core facility at the Cell Therapy Institute, PMU Salzburg. We thank Dr. Stefan Hainzl, EB House Austria, Department of Dermatology, University Hospital of the Paracelsus Medical University Salzburg, Austria, for the immortalization of primary human keratinocytes and fibroblasts. We further thank Ariane Benedetti for technical assistance and for performing the skin histology. In Seattle, we thank Kassidy Benoscek and Florence Roan for help in obtaining skin and blood samples, Thien Son-Nguyen for frozen PBMC, Alice Weidemann for help with CyTOF, Vivian Gersuk for help with RNA-seq, and Scott Presnell, Matt Dufort and Hannah DeBerg for helpful discussions on RNA-seq analysis.

## Data and materials availability

Raw and processed RNA-seq data can be found in Supplementary Data Excel Files 1 and 2. Immediately upon publication, RNA-seq data will be deposited in the gene expression omnibus (GEO), and CyTOF data will be made available at flowrepository.org

## Author contributions

M.M.K., P.A.M., B.H., S.R.V., T.D., D.J.C. and I.K.G. designed the experiments; M.M.K., M.D.R. and I.K.G. developed xenografting methods; M.M.K., P.A.M., B. H., S.R.V., S.M. and T.D. performed experiments; B.H. performed computational analysis; L.K-C. and E.G. performed analysis of TDL samples, M.R.B. planned and supervised the analysis of TDL samples, S.A.L. helped develop the CyTOF panels; G.B. helped set up the sort panels; D.J.C. and I.K.G. wrote the manuscript; M.M.K., P.A.M., B.H., S.R.V., S.M., T.D. and M.D.R. reviewed and edited the paper; D.J.C. and I.K.G. supervised the project.

---

**Fig. S1:**
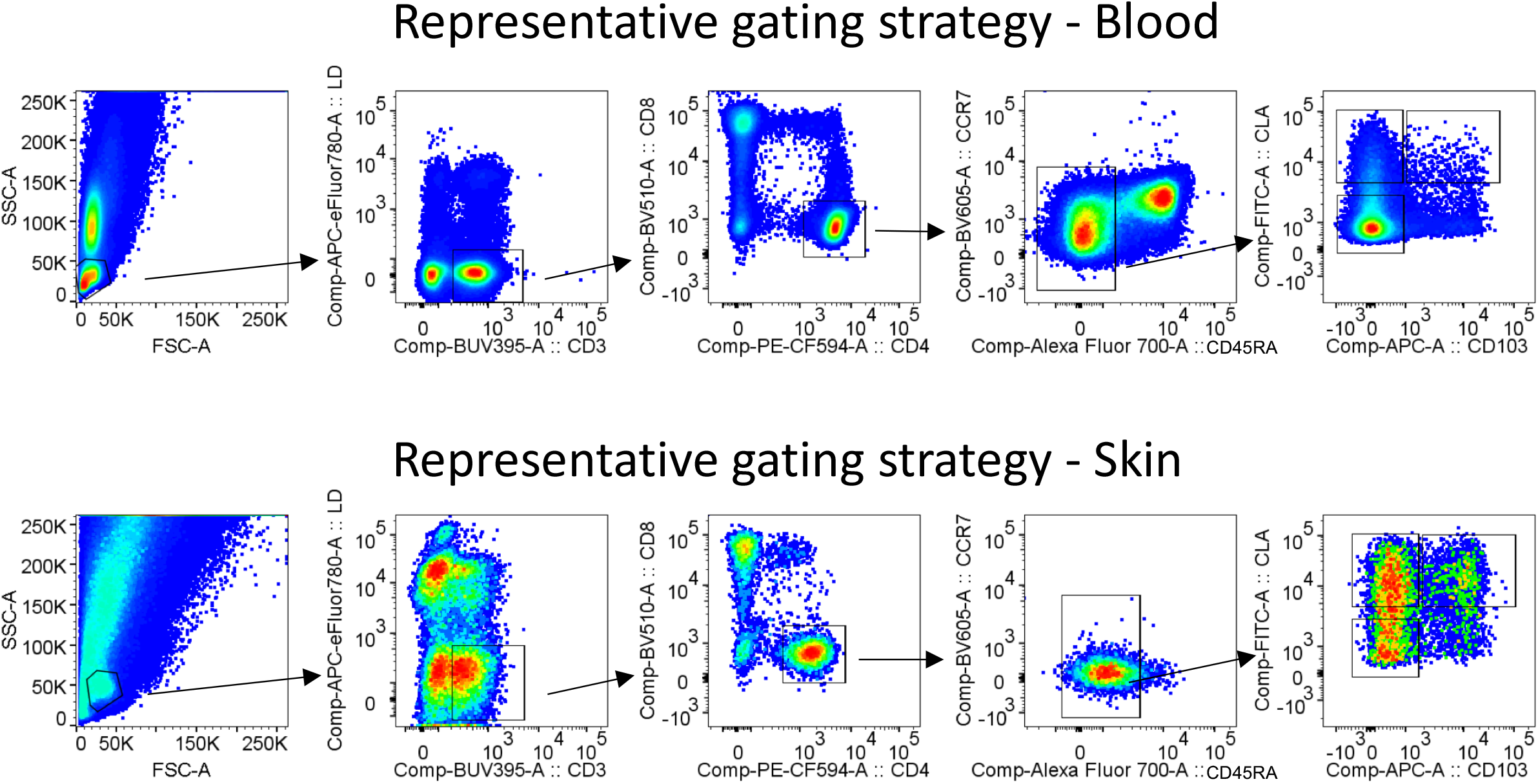
Representative flow cytometry gating strategies used to identify T cell subsets in human blood and skin.

**Fig. S2:**
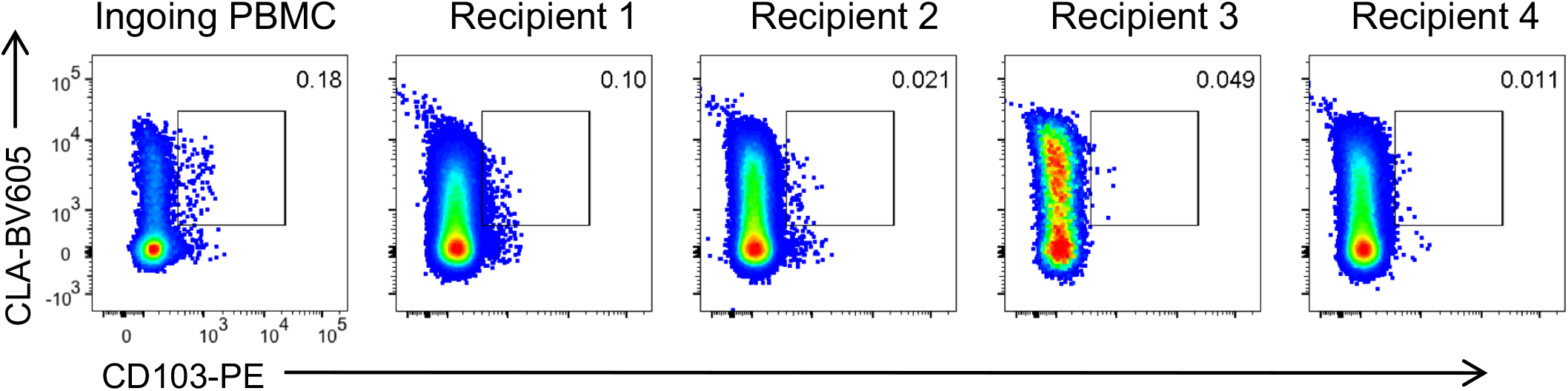
CD103 expression is not induced on human CD4^+^ T cells in NSG mice. 3×10^6^ PBMC were transferred into NSG mice and the spleens were analyzed by flow cytometry 49 days later. Representative flow cytometry analysis of CLA and CD103 expression by gated CD4^+^CD45^+^ T cells in the ingoing transferred PBMC and in cells recovered from the spleen of four individual recipients.

**Fig. S3:**
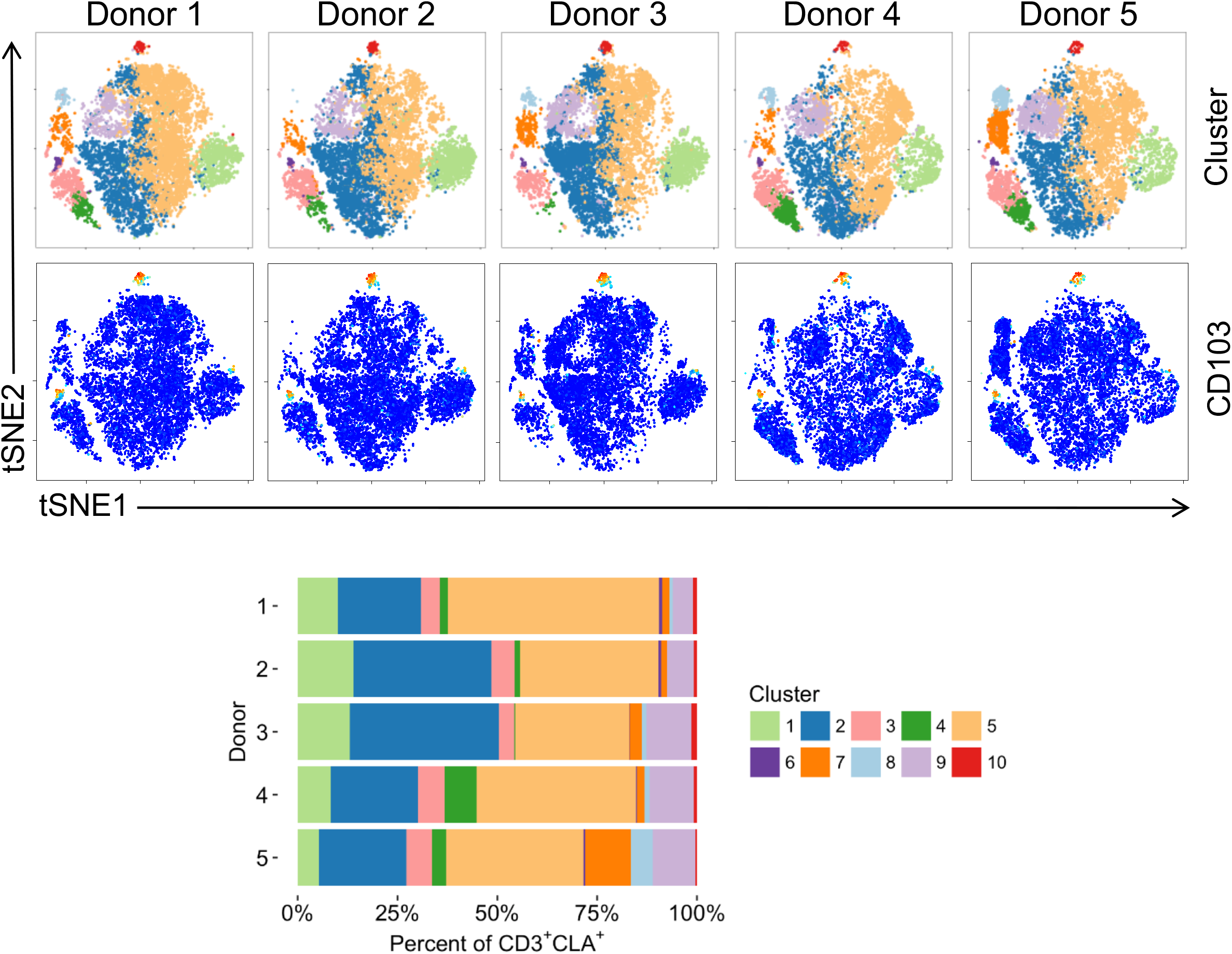
CyTOF analysis of CLA^+^ T cells in PBMC. (Top) tSNE analysis of gated CD3^+^CLA^+^ T cells from 5 individual donors showing clustering or CD103 expression as indicated. (Bottom) Stacked bar graph showing the frequency of cells in each of the 10 tSNE-defined clusters in each of the 5 donors examined.

**Fig. S4:**
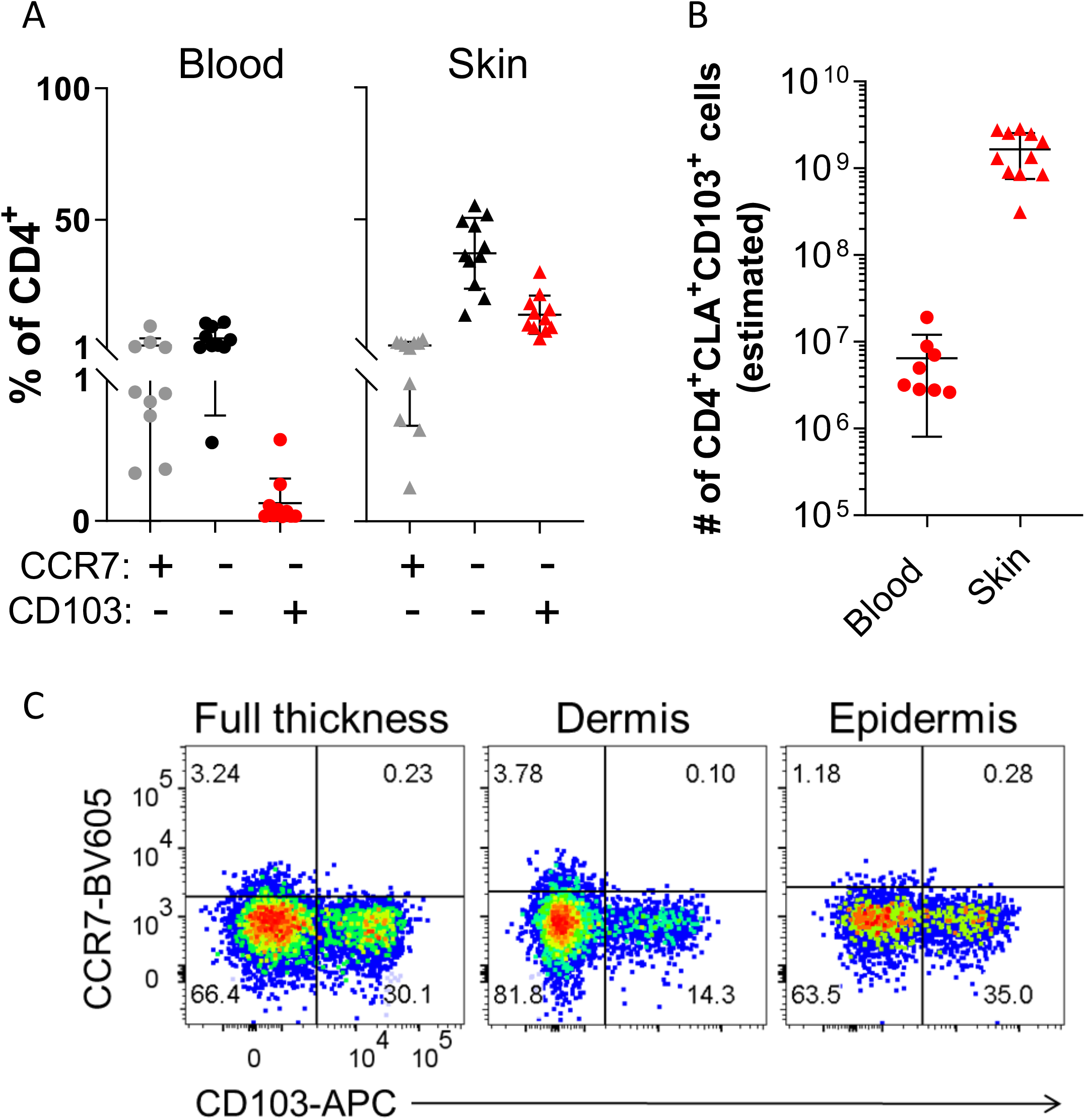
Frequencies of CLA^+^ T cell subsets in blood and skin. **(A)** Graphical summary of the proportions of CLA^+^CCR7^+^CD103^−^, CLA^+^CCR7^−^CDl03^−^, and CLA^+^CCR7^−^ CD103^+^ T cells among live gated CD4^+^CD45RA^−^T cells from blood and skin. **(B)** Estimation of CD4^+^CLA^+^CD103^+^ T cell numbers in skin and blood of different subjects were based on total T cell numbers in these tissues as determined in reference *(14),* the frequency of CD4^+^ T cells in each tissue as determined by the investigator’s labs, and the frequency of CD103^+^CLA^+^ T cells in each tissue as shown in part (A) of this figure. **(C)** Flow cytometric analysis of CCR7 and CD103 expression by gated CD4^+^CD45RA^−^CLA^+^ memory T cells from total full thickness skin (left), or from separated dermal (middle) or epidermal (right) compartments.

**Fig. S5:**
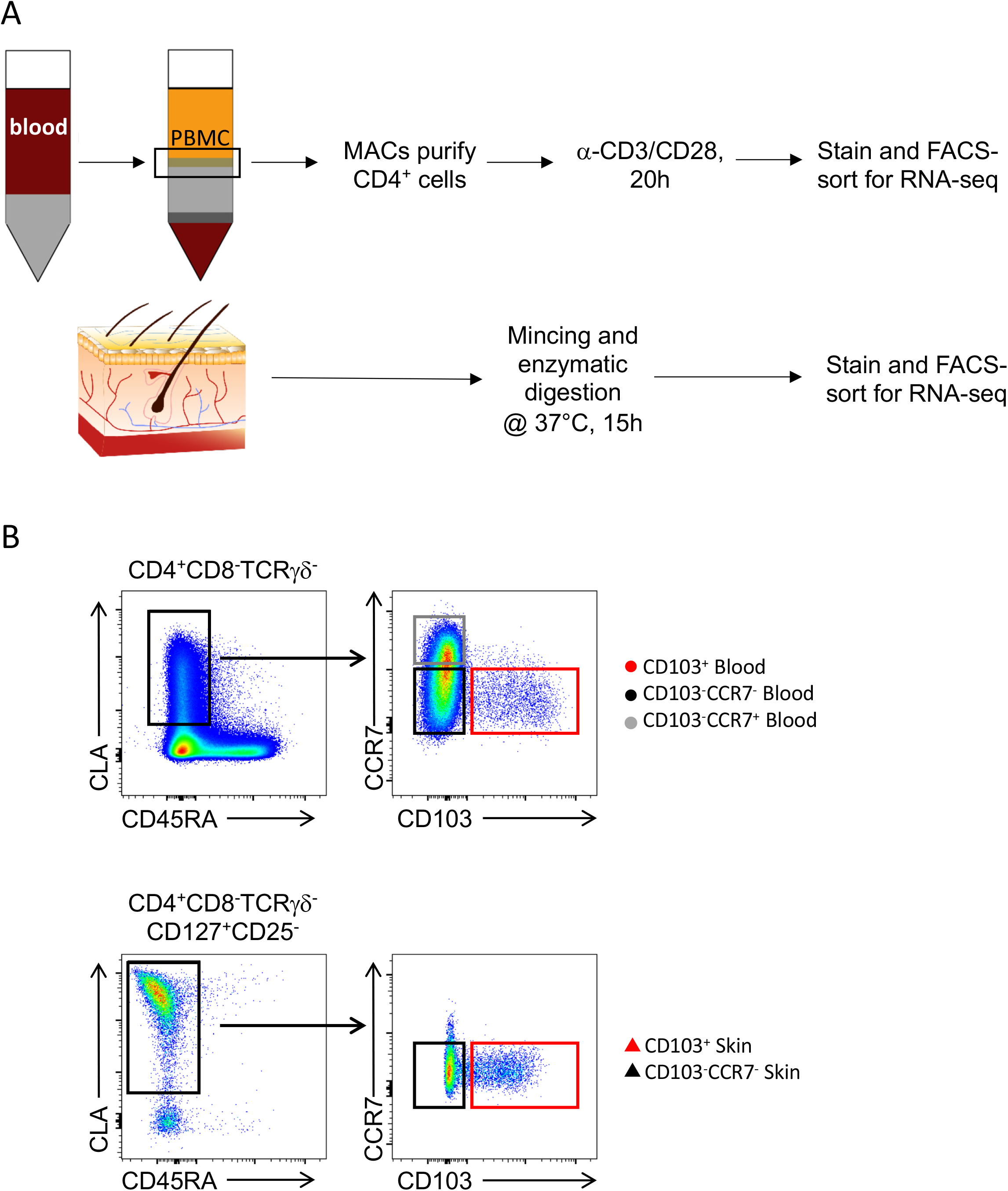
Experimental schematic of cell isolation and sort gates for RNA-seq. **(A)** Experimental schematic for processing and sorting of T cells from blood and skin for RNA-seq analysis. **(B)** Representative flow cytometry analysis of T cells isolated from blood or skin showing approximate gates used to sort the indicated populations for RNA-seq analysis.

**Fig. S6:**
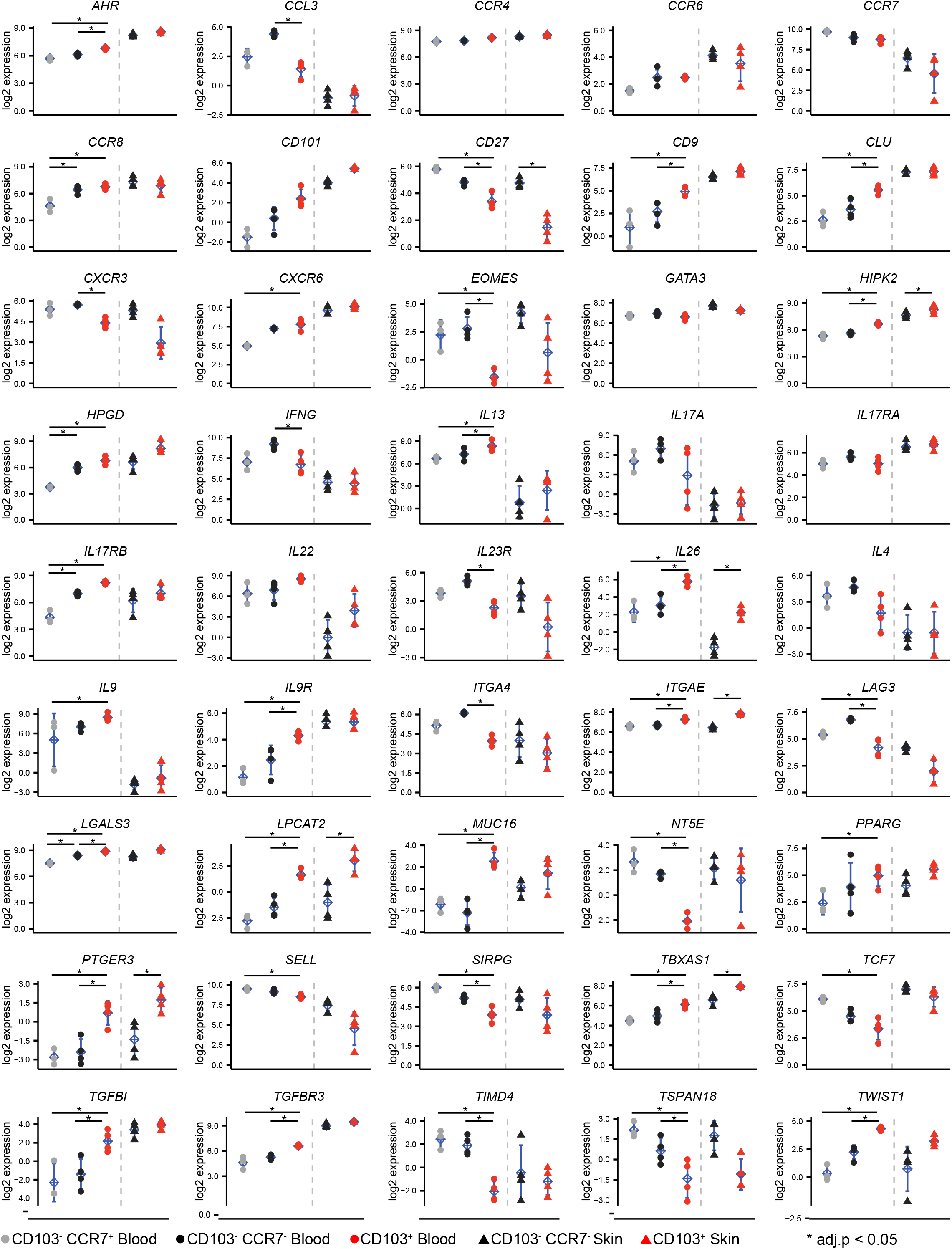
Expression of selected genes in sorted CD4^+^CLA^+^ T cells from blood and skin. Adjusted p.value computed by fitting a linear model with the displayed comparisons as contrasts.

**Fig. S7:**
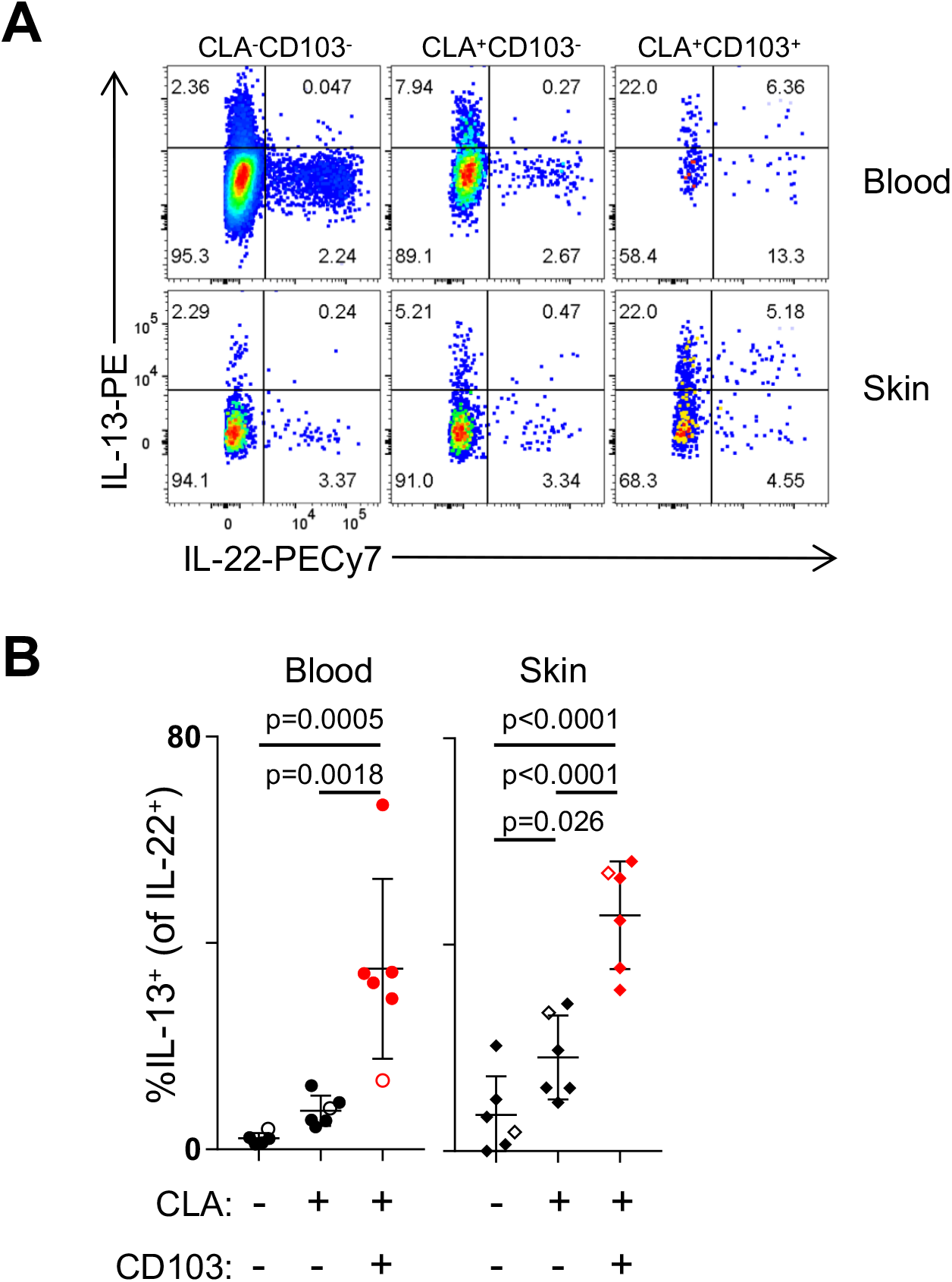
CLA^+^CD103^+^ T cells from human blood and skin coproduce IL-22 and IL-13. **(A)** Representative flow cytometry analysis of IL-22 and IL-13 production by the indicated populations of CD4^+^CD45RA^−^ T cells from paired blood and skin following PMA/ionomycin stimulation. **(B)** Graphical summary of the proportion of IL-13^+^ cells among IL-22^+^ CLA^−^ CD103^−^, CLA^+^CD103^−^ and CLA^+^CD103^+^ CD45RA^−^ memory T cells as indicated in blood and skin. Open symbols represent data from a subject with mammary carcinoma. Significance determined by one-way repeated measures ANOVA with Tukey’s post-test for pairwise comparisons.

**Fig. S8:**
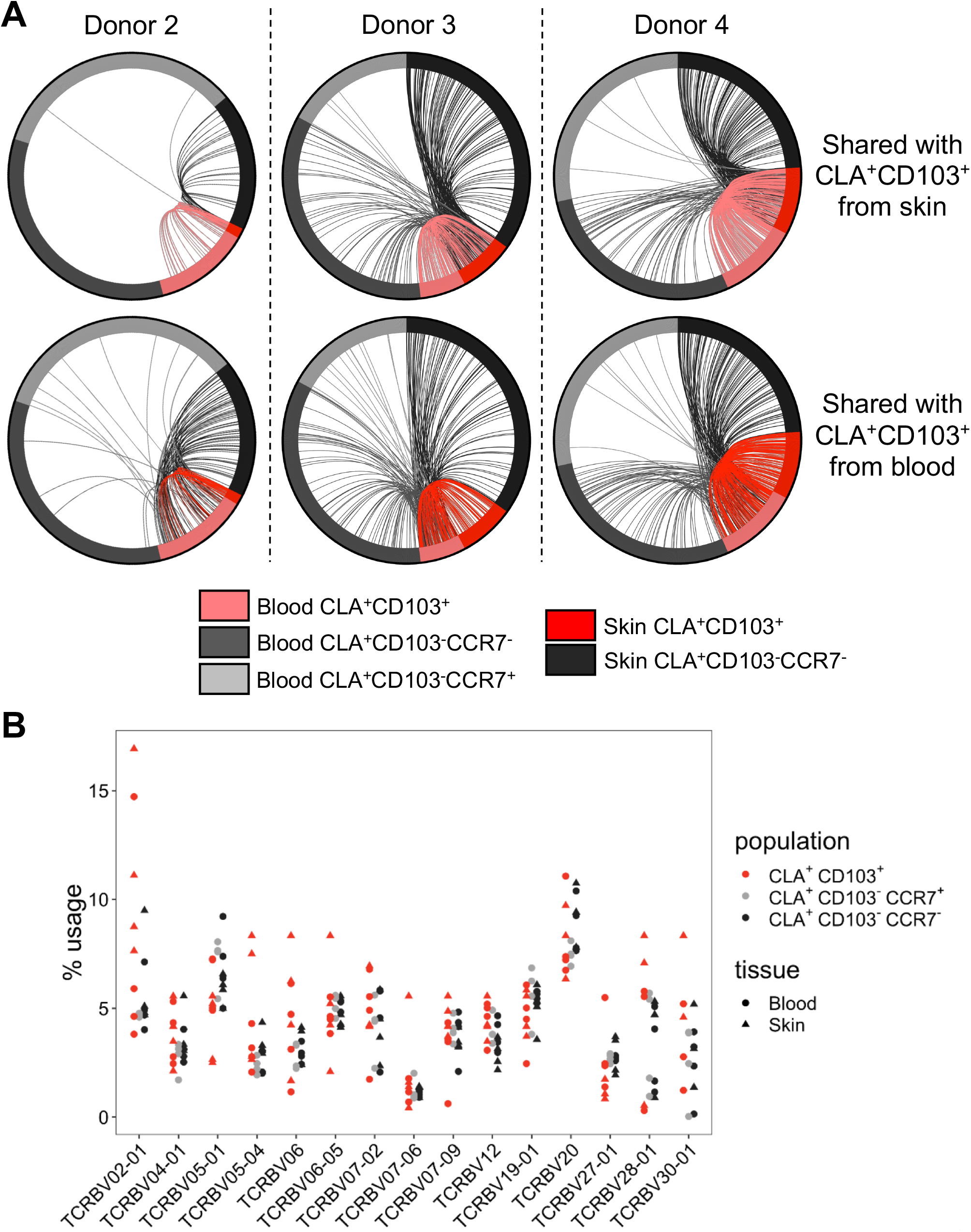
Analysis of TCRβ repertoire overlap and Vβ gene usage in CLA* T cells from blood and skin. **(A)** Circle plots of unique productive TCRβ sequences from each of the indicated populations of CLA^+^ memory T cells from each donor examined. Connections highlight sequences from blood or skin CLA^+^CD103^+^ cells found in each of the other populations as indicated. For visualization, sequences were downsampled (weighted for relative abundance) for populations containing >1000 unique sequences. **(B)** Graphical analysis of TCR Vβ segment usage in the indicated populations from across all four donors examined. The plot includes V gene segments that have a usage ≥5% in at least one sample.

**Fig. S9:**
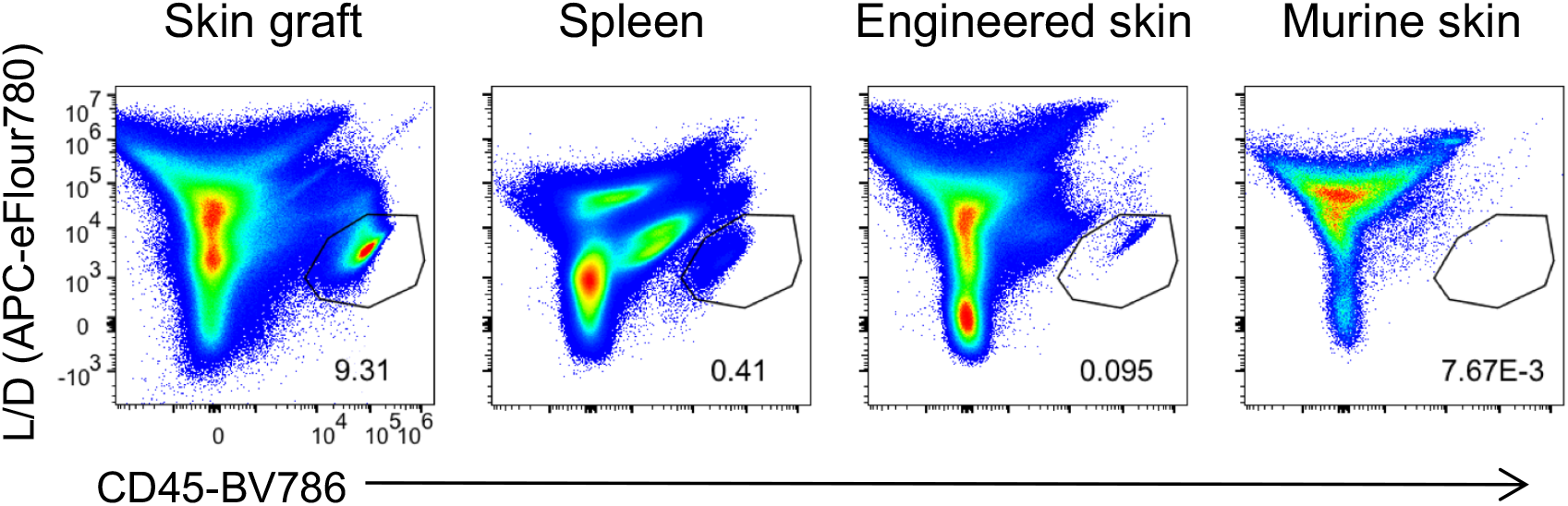
Human skin-derived cells do not infiltrate murine skin. Representative flow cytometry analysis of human CD45 expression versus live/dead stain of cells isolated from full-thickness skin graft, spleen, engineered skin, and adjacent murine skin 5 weeks after skin grafting.

**Table S1:** Detailed list of antibodies and reagents

**Tissue preparation**
| Reagent | Company | Catalog number |
| --- | --- | --- |
| Collagenase Type 4 | Worthington | LS004186 |
| DNase | Sigma-Aldrich | DN25 |
| RPMI 1640 | Gibco | 31870074 |
| human serum | Sigma-Aldrich | H5667/H4522 |
| Penicillin/streptavidin | Sigma-Aldrich | P0781 |
| L-Glutamine | Gibco | A2916801 |
| NEAA | Gibco | 11140035 |
| Sodium-Pyruvat | Sigma-Aldrich | S8636 |
| b-Mercaptoethanol | Gibco | 31350-010 |
| PBS | Gibco | 14190169 |
| Ficoll Paque Plus | GE-Healthcare | GE17-1440-02 |
| Dispase II | Sigma-Aldrich | D4693 |

**Cellular activation**
| Reagent | Company | Catalog number |
| --- | --- | --- |
| CD4 microbeads | Miltenyi | 130-045-101 |
| Immuno Cult Human CD3/CD28 T cell activator | Stemcell Technologies | 10971 |
| PMA | Sigma-Aldrich | P8139 |
| Ionomycin | Sigma-Aldrich | I06434 |
| Brefeldin A | Sigma-Aldrich | B6542 |
| Cytofix/Cytoperm kit | BD | RUO 554714 |
| Recombinant human IL-2 | Immunotools | 11340023 |

**Skin cell culture and transplantation**
| Reagent | Company | Catalog number |
| --- | --- | --- |
| Epilife | Gibco | MEPICF500 |
| DMEM | Gibco | 11960-044 |
| MEM | Gibco | 11380037 |
| TrypLE express | Gibco | 12604021 |
| Primocin | invitrogen | ant-pm-1 |
| Histoacryl | BRAUN | 1050044 |

**Tissue preparation from mice**
| Reagent | Company | Catalog number |
| --- | --- | --- |
| BD Pharm Lyse | BD | 555899 |
| Collagenase from Clostridium histolyticum | Sigma-Aldrich | C9407 |
| Hyaluronidase | Sigma-Aldrich | H3506 |

**CyTOF**
| Reagent | Company | Catalog number |
| --- | --- | --- |
| PBS (Ca and Mg free) | Sigma | D8537 |
| Cisplatin, 100 mM in DMSO | Enzo Life Sciences | ALX-400-040-M250 |
| Human TruStain FcX | BioLegend | 422302 |
| MaxPar Nuclear Antigen Staining Buffer Set | Fluidigm | 201063 |
| MaxPar Fix and Perm Solution | Fluidigm | 201067 |
| Cell-ID Intercalator-Ir, 125 uM (500X) | Fluidigm | 201192A |
| MaxPar EQ Four Element Calibration Beads | Fluidigm | 201078 |

**Histology**
| Reagent | Company | Catalog number |
| --- | --- | --- |
| TissueTek O.C.T. Compound | Sakura | TTEK |
| Hemalum solution acid acc. to Mayer | Carl Rorth | T865.1 |
| Eosin Y aqueous solution | Sigma | HT110232 |
| ProLong™ Gold Antifade Mountant with DAPI | Invitrogen | P36931 |

**Antibodies**
| Reagent | Company | Catalog number |
| --- | --- | --- |
| CLA Alexa Fluor 647 (HECA-452) | Biolegend | 321309 |
| CLA FITC (HECA-452) | Biolegend | 321306 |
| CLA bv605 (HECA-452) | BD | 563960 |
| CD103 APC (Ber-ACT8) | Biolegend | 350216 |
| CD103 FITC (B-Ly7) | eBioscience | 11-1038-42 |
| CD103 PE (Ber-ACT8) | Biolegend | 350206 |
| CD103 PE-Cy7 (Ber-ACT8) | Biolegend | 350212 |
| CD3 BV421 (UCHT1) | Biolegend | 300434 |
| CD3 BUV805 (UCHT1) | BD | 565515 |
| CD3 PE (HIT3a) | BD | 561803 |
| CD3 PE-Cy5.5 (UCHT1) | eBioscience | 15-0038-42 |
| CD4 Brilliant Blue 700 (SK3) | BD | 566392 |
| CD4 PE-594 (RPA-T4) | Biolegend | 300548 |
| CD4 PE-Cy5 (RPA-T4) | BioLegend | 300510 |
| CD4 Alexa Fluor 700 (RPA-T4) | Biolegend | 300526 |
| CD8 Pacific Orange (3B5) | eBioscience | MHCD0830 |
| CD8 bv510 (RPA-T8) | Biolegend | 301048 |
| CD8 bv785 (RPA-T8) | BioLegend | 301046 |
| CD8 BUV496 (RPA-T8) | BD | 564804 |
| CD9 PE (M-L13) | BD | 341647 |
| CD25 PE-Cy7 (BC96) | eBioscience | 302612 |
| CD27 PE-594 (M-T271) | BioLegend | 356422 |
| CD27 BV570 (O323) | Biolegend | 302825 |
| CD27 BV711 (M-T271) | BioLegend | 356430 |
| CD45 BV785 (HI30) | Biolegend | 304048 |
| CD45RA Alexa Fluor 700 (HI100) | Biolegend | 304120 |
| CD69 BV605 (FN50) | Biolegend | 310938 |
| CD69 PE (FN50) | BD | 555531 |
| CD101 PerCP-Cy5.5 (BB27) | Biolegend | 244438 |
| CD127 PE-Cy7 (A019D5) | BioLegend | 351320 |
| CD127 bv 421 (HIL-7R-M21) | BD | 560823 |
| CCR4 PerCP-Cy5.5 (L291H4) | Biolegend | 359406 |
| CCR4 PE-CF594 (1G1) | BD | 565391 |
| CCR8 PE (L263G8) | Biolegend | 360604 |
| CD45RA BUV563 (HI100) | BD | 565702 |
| CD45RA PE-e610 (HI100) | eBioscience | 61-0458-42 |
| CCR6 PE-Cy7 (R6H1) | eBioscience | 25-1969-42 |
| CXCR3 bv421 (1C6/CXCR3) | BD | 562558 |
| CCR6 PE-Cy7 (G034E3) | Biolegend | 353418 |
| CCR6 BV605 (G034E3) | Biolegend | 353420 |
| CCR7 APC-Cy7 (G043H7) | Biolegend | 353211 |
| CCR7 bv421 (G043H7) | BioLegend | 353232 |
| CCR7 bv605 (G043H7) | Biolegend | 353224 |
| IFN-g BV605 (4S.B3) | BD | 563731 |
| IL-17A PerCP-Cy5.5 (eBio64DEC17) | eBioscience | 45-7179-42 |
| IL-13 PE (JES10-5A2) | BD | 554571 |
| IL-22 PE-Cy7 (22URTI) | eBioscience | 25-7229-42 |
| GM-CSF bv421 (BVD2-21C11) | BD | 562930 |
| Fixable Viability Dye eFluor® 780 | eBioscience | 65-0865-14 |
| LIVE/DEAD Fixable Aqua Dead Cell Stain Kit | ThermoFisher Scientific | L34957 |
| TCRgd PerCP-eFluor710 (B1.1) | eBioscience | 46-9959 |
| IF primary antibody: rabbit anti-human NC-1 domain of type VII collagen (LH7.2) | kindly provided by Dr. Alexander Nyström,<br>University of Freiburg, Germany |  |
| IF secondary antibody: goat anti-rabbit A488 | ThermoFisher | A11008 |
| mLy6G (Gr-1) InVivoMab RB6-8C5 | BioXcell | BE0075 |

Antibodies for CyTOF
| Reagent | Company | Catalog number |
| --- | --- | --- |
| CLA FITC (HECA-452) | BioLegend | 321306 |
| Integrin-beta7 APC (FIB504) | BioLegend | 321208 |
| CTLA4 147Sm (14D3) | Fluidigm | Custom |
| Ki-67 161Dy (B56) | Fluidigm | 3161007B |
| Foxp3 162Dy (259D/C7) | Fluidigm | 3162024A |
| CCR6 141Pr (G034E3) | Fluidigm | 3176022A |
| CD19 142Nd (HIB19) | Fluidigm | 3142001B |
| CD123 143Nd (6H6) | Fluidigm | 3143014B |
| CD38 144Nd (HIT2) | Fluidigm | 3144014B |
| CD4 145Nd (RPA-T4) | Fluidigm | 3145001B |
| IgD 146Nd (IA6-2) | Fluidigm | 3146005B |
| CD16 148Nd (3G8) | Fluidigm | 3148004B |
| CCR4 149Sm (205410) | Fluidigm | 3149029A |
| CD103 151Eu (Ber-ACT8) | Fluidigm | 3151011B |
| CD45RO 152Sm (UCHL1) | Fluidigm | Custom |
| CD45RA 153Eu (HI100) | Fluidigm | 3153001B |
| CD45 154Sm (HI30) | Fluidigm | 3154001B |
| CD27 155Gd (M-T271) | Fluidigm | 3155001B |
| CXCR3 156Gd (1C6) | Fluidigm | 3156004B |
| CD10 158Gd (HI10a) | Fluidigm | 3158011B |
| CD161 159Tb (HP-3G10) | Fluidigm | 3159004B |
| CD14 160Gd (M5E2) | Fluidigm | 3160001B |
| Anti-APC 163Dy | Fluidigm | 3163001B |
| CCR0 164Dy (314305) | Fluidigm | Custom |
| CD127 165Ho (A019D5) | Fluidigm | 3165008B |
| CD24 166Er (ML5) | Fluidigm | 3166007B |
| CCR7 167Er (150503) | Fluidigm | 3167009A |
| CD8 168Er (SK1) | Fluidigm | 3168002B |
| CD25 169Tm (M-A251) | Fluidigm | 3169003B |
| CD3 170Er (UCHT1) | Fluidigm | 3170001B |
| IgM 172Yb (MHM-88) | Fluidigm | 3172004B |
| HLA-DR 173Yb (L243) | Fluidigm | 3173005B |
| Anti-FITC 174Yb | Fluidigm | 3174006B |
| CRTh2 175Lu (BM16) | Fluidigm | Custom |
| CD56 176Yb (HCD56) | Fluidigm | 3176001B |

